# Sleep deprivation differentially recruits amygdala subregions in male and female mice

**DOI:** 10.64898/2026.11.09.765501

**Authors:** Nikita K Koziel Ly, Chia-Luen Leu, Daniel D Lam, Violetta Molokopoy, Rhîannan H Williams, Melissa J Chee

## Abstract

Sleep deprivation (SD) acutely increases negative affect, and chronic insufficient sleep is associated with an increased risk of depression. Since the amygdala regulates both emotional processing and sleep, and SD dysregulates amygdalar function, amygdala functionality may link sleep loss to emotional outcomes. To better understand this, the specific sleep-impacted amygdalar nuclei need to be fully defined. To determine the impact of acute sleep loss on the relative activation of specific amygdalar regions, we screened for the neuronal activity marker c- Fos in wildtype mice that slept undisturbed, underwent 4-h SD, or 4-h SD with 2-h recovery sleep (RS). Application of whole-brain clearing and automated cell counting identified increased c-Fos expression within the cortical subplate of both male and female mice that was localized to amygdalar regions. As the amygdala comprises multiple distinct subnuclei, we parcellated and quantified c-Fos expression within each amygdalar subnucleus. Interestingly, we found that SD-induced differential c-Fos expression patterns between sexes. In male mice, SD increased c-Fos expression across several amygdalar subnuclei, which largely reversed with RS. By contrast, SD in females increased c-Fos expression only in the anterior basolateral amygdala and the intercalated amygdala, and both subnuclei remained activated even with RS. These findings highlight sex-specific spatial distribution of SD-activated amygdala cells, which may contribute to differences in emotional processing following sleep loss and offer potential anatomical targets for mitigating SD-related mood deficits.

**Highlights:**

- An unbiased screen, using c-Fos as a measure of neuronal activity highlighted amygdalar regions within the cortical subplate as structures activated by sleep deprivation (SD).
- SD broadly increased amygdalar c-Fos in male mice but produced a more restricted anatomical pattern in females.
- The anterior basolateral amygdala (BLAa) and intercalated amygdala (IA) were activated by SD in both sexes.

## 1 Introduction

Adults are advised to obtain 7–9 hours of sleep per night for optimal health (Watson et al., 2015), yet one in four adults aged 18–79 regularly sleeps less than the daily recommendation (Canada, 2019). Acute sleep loss prominently amplifies negative mood states, including anxiety (Pires et al., 2016; Short & Louca, 2015), irritability (Dement, 1960), and anger (Minkel et al., 2012). Furthermore, chronic sleep deprivation (SD) may increase the risk for mood disorders such as depression (Roberts & Duong, 2014), particularly in women (Conklin et al., 2018).

The brainstem, thalamus, hypothalamus, and basal forebrain are well-established regulators of sleep-wake balance (Brown et al., 2012; Saper et al., 2010). However, these sleep-wake circuits alone are unlikely to explain the mood deficits associated with acute or chronic SD. Indeed, human imaging studies indicate that additional brain regions, beyond those that regulate sleep-wake balance, are differentially activated with sleep loss. For example, a meta-analysis of 23 resting state functional magnetic resonance imaging studies reported increased striatal activity and decreased medial frontal gyrus activity in healthy participants after acute SD (Zhang et al., 2024).

Cortical regions are strongly connected to limbic structures such as the amygdala, a key node for mood and emotional processing, and functional neuroimaging shows SD alters connectivity between the cortex and amygdala (Yoo et al., 2007). Specifically, SD reduces amygdala connectivity within the dorsolateral prefrontal and anterior cingulate cortices (Shao et al., 2014), which support executive control and emotional regulation, respectively. In contrast, stronger amygdala–anterior cingulate connectivity is associated with better mood in healthy participants and patients with depression (Chai et al., 2023).

In addition to mood-related functions, the amygdala is also implicated in sleep. For example, human imaging studies have recorded increased amygdalar blood flow and glucose metabolism during rapid eye movement sleep (REM; Corsi-Cabrera et al., 2016; Maquet et al., 1996; Nofzinger et al., 1997). Moreover, increasing sleep pressure during wake is related to increased amygdalar reactivity while increasing sleep opportunity suppresses amygdalar reactivity (van der Helm et al., 2011; Wassing et al., 2019). These studies suggested heightened amygdalar activity during sleep or as sleep need accumulates.

Causal evidence comes from preclinical work, in which amygdalar stimulation improved recovery from REM-specific sleep loss (Smith & Miskiman, 1975). However, these effects are not uniform across the amygdala, as intervention studies in rodents indicate that distinct subnuclei contribute to different aspects of sleep regulation. Basolateral amygdala (BLA) lesions increase non-rapid eye movement sleep (NREM; Zhu et al., 1998), and subpopulations of BLA neurons are active during recovery sleep (Bocchio et al., 2016). Recent datasets show that specific changes in amygdalar cell populations drive different state outcomes. For example, neurotransmitter fluctuations within the BLA triggers REM (Hasegawa et al., 2022, 2024), inactivation of the central amygdala (CEA) decreases REM (Sanford et al., 2002, 2006; Tang et al., 2005), and inhibitory CEA-neurotensin cells promote and maintain NREM (Ma et al., 2019). Taken together, amygdala circuits are thus poised to contribute to sleep loss-related emotional disturbances, but the specific amygdalar nuclei activated during SD are not yet well-defined.

To better define engagement of amygdalar subnuclei with relevance to sleep loss and affective processing, we performed an unbiased screen for c-Fos expression throughout top-level structures in the intact brain. This provided a snapshot of inferred cellular activity, as marked by a thresholded c-Fos value. We observed that SD increased activity within the cortical subplate, particularly in amygdalar regions of male and female wildtype mice. Next, we quantified and mapped the distribution of c-Fos expression within each amygdalar subnucleus. Interestingly, there was a notable sex difference in the amygdalar response to SD. In males, SD increased c-Fos expression across several amygdalar subnuclei, which largely reversed with recovery sleep (RS). By contrast, in females, SD increased c-Fos expression only in the anterior part of the basolateral amygdalar nucleus (BLAa) and the intercalated amygdalar nucleus (IA), and both subnuclei remained activated despite sleep opportunity.

Together, these findings point to specific recruitment of the amygdala during and after sleep loss. Importantly, our work highlights the unique spatial distribution of SD-activated cells within individual amygdala subnuclei that provide candidates for future studies of SD-related affective changes.

## 2 Materials and Methods

### 2.1 Animals

All procedures were approved by the responsible authority of the Regierung von Oberbayern (Government of Upper Bavaria; ROB-55.2-2532.Vet_02-17-162) and the Animal Care Committee at Carleton University (Canada). C57BL/6JOlaHsd wildtype mice (Harlan/Envigo) were housed on a 12:12 h light-dark cycle at ambient temperature (21–24°C) and given *ad libitum* access to water and standard rodent chow (Altromin 1328, Lage, Germany; Teklad Global Diets 2014, Envigo, Mississauga, Canada. Protocol parameters, enrichment, and animal handling were unified across sites.

### 2.2 Sleep protocol

Mice were sleep deprived for 4 h from the start of the light cycle at Zeitgeber time (ZT) 0 to ZT4. Mice undergoing sleep deprivation (SD) remained in their homecage and were visually monitored for signs of sleep. When mice began to close their eyes, were quiescent, or appeared hunched, their cage environment was disturbed by moving their nest (e.g., plastic huts, cardboard houses, nestlet) and/or their vibrissae or fur was gently stroked with a paintbrush.

Mice in the recovery sleep (RS) group underwent SD for 4 h, as described above, and then were allowed to sleep undisturbed for 2 h. All bedding, housing, and nestlets were returned before the end of the SD period to facilitate sleep onset and RS.

Corresponding time-matched controls were left undisturbed in their homecage until sacrifice at either ZT4 for the SD group or ZT6 for the RS group. These control mice were housed separately to mice experiencing SD.

### 2.3 Antibody characterization

The primary antibodies used for iDISCO+ and immunohistochemistry are listed in **Table 1**

**Table 1.** List and details of primary antibodies.

| Antibody | Immunogen | Clonality,<br>Isotype | Source,<br>Catalogue no.,<br>Lot no. | RRID | Titer |
|---|---|---|---|---|---|
| Rabbit anti c-Fos | Amino acid 2–17 rat FOS | Polyclonal,<br>IgG | Synaptic Systems,<br>226003,<br>N/A | AB_2231974 | 1:2,000 |
| Rabbit anti-c-Fos | Full length human FOS | Polyclonal,<br>IgG | Abcam,<br>ab190289,<br>1091820-1 | AB_2737414 | 1:2,000 |

Rabbit anti-c-Fos antibody (Synaptic Systems, Göttingen, Germany) was raised against rat synthetic peptide MFSGFNADYEASSSR and recognizes amino acids 2–17 of rat c-Fos protein. Specificity of this antibody was validated using Western blot, which showed increased band strength in HeLa cell extracts after stimulation with 12-O-tetradecanoylphorbol-13-acetate (Synaptic Systems, manufacturer correspondence).

Rabbit anti-c-Fos antibody (Abcam, Cambridge, UK) recognizes full length human FOS. It is a 41 kDa protein that has been confirmed by Western blot. Specificity was also determined *via* immunostaining in HeLa cells stimulated with 20% fetal bovine serum, which revealed c-Fos labelling in only stimulated cells compared to unstimulated cells treated with phosphate buffered saline (Abcam, manufacturer correspondence).

The secondary antibodies are listed in **Table 2**. Each was raised in goat or donkey against the host species of the corresponding primary antibody.

**Table 2.**
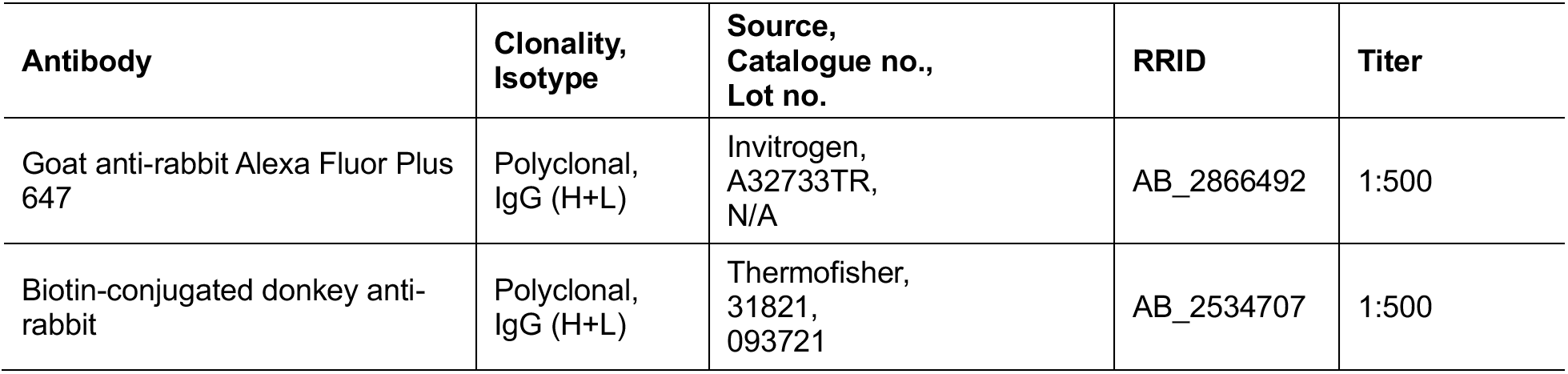
List and details of secondary antibodies.

### 2.4 Immunolabeling-enabled three-dimensional imaging of solvent-cleared organs clearing (iDISCO+)-based whole-brain clearing

Brains were prepared for whole brain imaging as previously described (Ertürk et al., 2012; Renier et al., 2016). To account for potential differences in batch processing, animals from each condition were split equally across processing days and conditions remained blinded.

#### Tissue fixation

Male and female mice (4–7 months; *n* = 4 per group) were treated with oral Melosus (1–2.5 mg/kg). Two hours later, mice were euthanized with a rising gradient of carbon dioxide and transcardially perfused with 0.01 M phosphate buffered saline (PBS) followed by 4% paraformaldehyde (PFA; in 0.01 M PBS). Brains were extracted from the skull and post-fixed overnight in 4% PFA at 4°C on a shaker and then transferred to 0.2% sodium azide in PBS at 4°C until ready for iDISCO+ processing.

#### Pretreatment

All incubation and washing steps occurred using individual glass jars (7–8 mL) placed on a rotating or orbital shaker to aid tissue penetration while averting bubble formation. Whole brains were washed in PBS three times for 30 min each at room temperature (RT), then were dehydrated in ascending concentrations of methanol (20%, 40%, 60%, 80%, and 100%) for 1 h each. Brains were washed twice in 100% methanol for 30 min each at RT, then for 1 h at 4°C. Brains were then bleached overnight in chilled 5% hydrogen peroxide in methanol (4°C).

The next day, brains were returned to RT then were rehydrated in descending concentrations of methanol (80%, 60%, 40%, and 20%) for 1 h each (RT). Brains were then washed twice for 1 h each in PBS with 0.2% Triton X-100 (PBT) at RT.

#### Immunolabeling

Brains were incubated in PBT with 20% dimethyl sulfoxide (DMSO) and 0.3 M glycine for 3–4 days at 37°C, then blocked in 6% normal goat serum (NGS; Life Technologies) prepared in PBT with 10% DMSO for 3–4 days at 37°C. Brains were incubated in rabbit anti-c-Fos primary antibody (1:2,000, Synaptic Systems; **Table 1**) in PBT with heparin (10 µg/mL; PBT-heparin), 5% DMSO, and 3% NGS for 7 days at 37°C. After 4–5 exchanges in PBT- heparin over 24 h, brains were incubated in goat anti-rabbit-conjugated Alexa Fluor Plus 647 (1:500; **Table 2**) for 7 days at 37°C, followed by another 4–5 exchanges in PBT-heparin over 24 h.

#### Tissue clearing

Brains were dehydrated in ascending concentrations of methanol (20%, 40%, 60%, 80%, and 100%) for 1 h each, then left in fresh 100% methanol overnight at RT. The next day, brains were incubated in a solution of 66% dichloromethane (DCM) and 33% methanol for 3 h at RT, then washed twice in 100% DCM for 15 min each. Finally, brains were incubated in dibenzyl ether for 4–5 days at RT prior to imaging.

### 2.5 Dual immunoperoxidase IHC and Nissl staining

#### Tissue processing

Male and female mice (4 months; *N* = 3–4 per group) were treated with Melosus (1–2.5 mg/kg) before they were euthanized with a rising gradient of carbon dioxide and transcardially perfused with 0.01 M PBS followed by 4% PFA. Brains were extracted from the skull and post-fixed overnight in PFA before being cryoprotected in PBS containing 30% sucrose and 0.05% sodium azide for at least 12 h at 4 °C.

All brains were sliced into six series of 30 μm-thick coronal sections using a freezing microtome (Leica SM2010, Wetzlar, Germany). Brain series were stored in anti-freeze comprising 50% formalin, 20% glycerol, 30% ethylene glycol at −20 °C until required for immunolabeling.

#### Immunoperoxidase immunohistochemistry (IHC)

As previously described (Bono et al., 2025; Chee et al., 2013), free floating brain slices were washed six times in PBS (RT) for 5 min each, then incubated in PBS containing 0.3% hydrogen peroxide for 30 min (RT) to reduce endogenous peroxidase activity. After another six 5-min PBS washes, slices were blocked in 3% normal donkey serum (NDS; Jackson ImmunoResearch Inc., West Grove, PA) prepared in PBT with 0.05% sodium azide (PBT-azide) for 2 h at RT, then incubated in rabbit anti-c-Fos primary antibody (1:2,000, Abcam; **Table 1**) for 16 h at RT. After six 5-min PBS washes at RT, the slices were incubated in donkey anti-rabbit biotinylated secondary antibody (1:500; **Table 2**) added to 3% NDS blocking solution in PBT for 1 h (RT) and then washed with three 10-min PBS washes. Slices were then treated with avidin-biotin peroxidase complex (ABC; 1:500; PK-6100, Vector Laboratories, Newark, CA) for 1 h (RT) in PBS, followed by three 10-min PBS washes. ABC was prepared minimum 30 min before use. Next, the slices were reacted with 3,3′-diaminobenzidine enhanced with nickel (SK-4100, Vector Laboratories) and monitored closely for black colour transformation (∼3 min). The reaction was terminated in distilled water then by washing in PBS three times for 10 min each (RT). The stained slices were mounted on Superfrost Plus microscope slides (Fisher Scientific, Ottawa, Canada), air-dried at RT, dehydrated in ascending concentrations of ethanol (50%, 70%, 95%, and 100%) for 3 min each, and cleared with xylene for a minimum of 2 h.

#### Nissl staining

Next, mounted and stained slices underwent Nissl-staining as previously described (Bono et al., 2022; Miller et al., 2024). Briefly, slides were dehydrated in ascending concentrations of ethanol (50%, 70%, 95%, and 100% × 3) for 3 min each (RT), delipidated in two exchanges of xylene (Millipore Sigma, Burlington, MA) for 10 min then 15 min, and rehydrated in descending concentrations of ethanol (100%, 95%, 70%, and 50%) and distilled water for 3 min each. To visualize cytoarchitectural boundaries, slides were submerged in 0.25% thionin (Millipore Sigma; pH 4.5) for maximum 5 s followed by three exchanges in distilled water. Finally, slides were dehydrated in ethanol and cleared in xylene as above, then coverslipped with Richard-Allan Scientific Mounting Media (Fisher Scientific) and 1.5 thickness glass (Fisher Scientific).

### 2.6 Microscopy

#### 2.6.1 Light sheet imaging

Cleared whole-brain samples were imaged through the horizontal plane on a light sheet fluorescence microscope (LaVision Ultramicrosope; Olympus MVX10 with objective lenses MV PLAPO 2XC with ImSpector Pro 64; LaVision BioTec, Göttingen, Germany; Analytical Pathology, Helmholtz Zentrum München). The ventral side of each brain was mounted in a holder and secured in place with two spikes penetrating the preoptic and hypothalamic regions. The holder was then placed in the imaging chamber filled with dibenzyl ether. Whole-brain imaging was conducted using a ×0.63 objective lens (0.1 numerical aperture) at 95% laser power (NKT Photonics, Birkerød, Denmark) and a scanning step-size of 7 µm. Wavelengths used: 520/40 nm excitation and 584/40 nm emission for autofluorescence; 640/30 nm excitation, and 690/50 nm emission for c-Fos imaging. A resolution of 2048×2048 pixels and 16-bit grayscale was applied for each horizontal 2D image.

#### 2.6.2 Brightfield imaging

As previously described (Chee et al., 2015), large field-of-view photomicrographs of whole brain sections that underwent dual IHC and Nissl-staining were acquired *via* a Plan Apochromat ×10 objective lens (0.45 numerical aperture) mounted on a Nikon Eclipse Ti2 inverted microscope (Nikon Instruments Inc., Mississauga, Canada) equipped with a fully motorized stage and DS-Ri2 color camera (Nikon). Brightfield images were captured with NIS-Elements (Nikon).

Acquired images were stitched with 10% overlap.

### 2.7 Image analysis

#### 2.7.1 Brain atlas registration and automated cell counting

To select an appropriate reference atlas, autofluorescence volumes from two cleared brains were registered to three volumetric mouse brain atlases: Perens et al. (2021; **Figure 1A**), Wang et al. (2020; **Figure 1B**), and (Young et al. 2021; **Figure 1C**). Registration quality was assessed by blinded visual inspection of atlas–sample overlap and by Pearson correlation of fluorescence intensities at corresponding voxel coordinates. The Perens atlas, which was optimized for iDISCO+ and light-sheet fluorescence microscopy data, showed the best correspondence with our samples (**Figure 1**) and was therefore used for all subsequent registrations and regional assignments. Registration was not adjusted for sex or substrain because suitable reference datasets, particularly for female brains, were unavailable.

**Figure 1.**
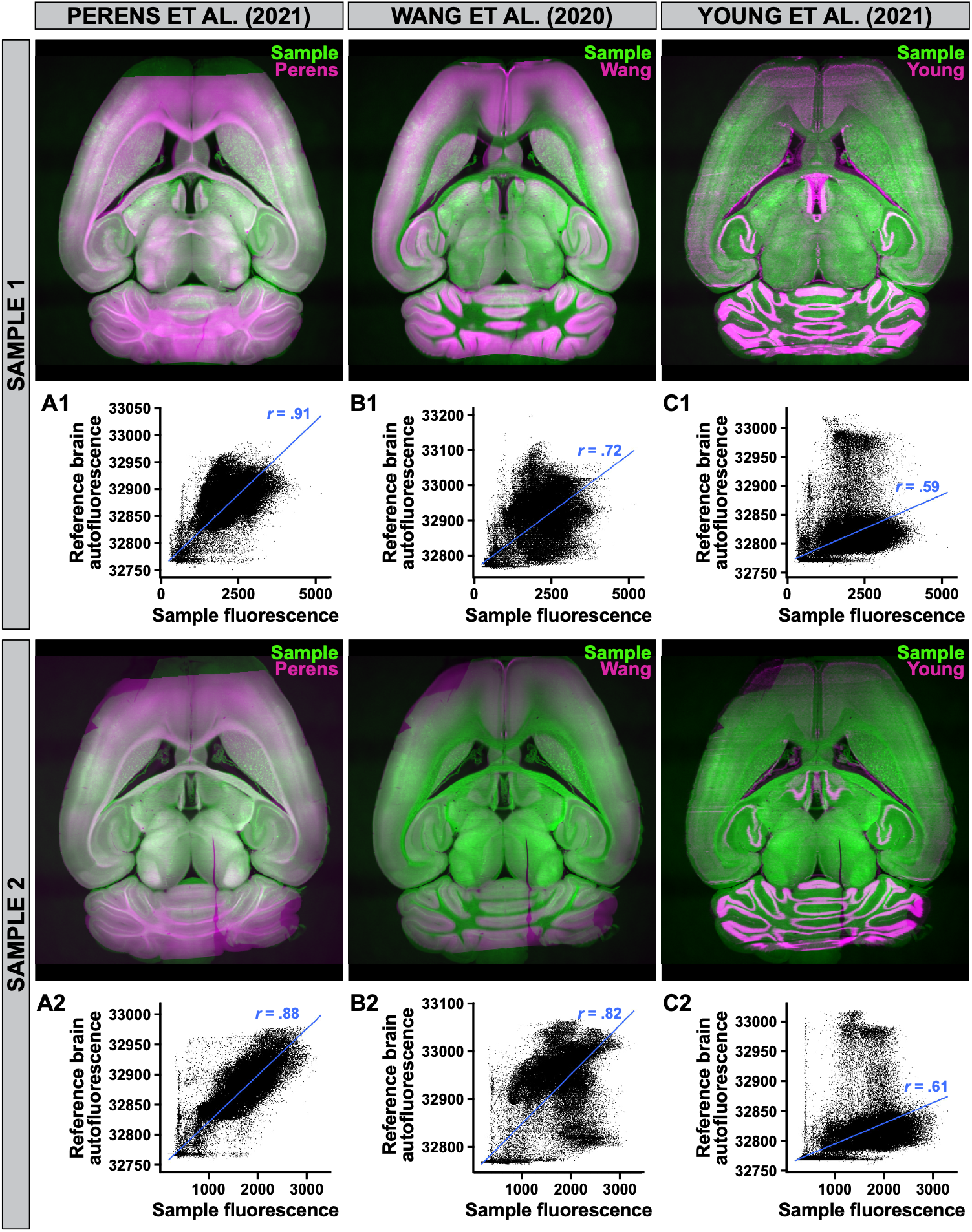
Registration of cleared brains to three mouse brain atlases. Pearson correlation of sample-to-atlas overlap of autofluorescence intensities at corresponding voxel coordinates from two whole brain samples (**1**, **2**) to three-dimensional mouse brain atlases by Perens et al. (2021; **A**), Wang et al. (2020; **B**), and Young et al. (2021; **C**). Overlap (white) determined from overlay of autofluorescence from the sample (green) and reference atlas (magenta) (top row).

Each c-Fos immunofluorescence volume was registered to the Perens atlas and processed with the Linux-based version of ClearMap (Renier et al., 2016; **Figure 2A–C**). Detection thresholds were defined separately for each batch and experimental group after inspection of three horizontal brain regions and determination of the maximum whole-brain fluorescence intensity. To accommodate within-batch variation while limiting threshold drift, sample-specific thresholds were constrained to within ±0.25 standard deviations of the group mean intensity. All thresholds were independently verified by a second examiner who was blinded to experimental condition. Light sheet photomicrographs, from one cortical region of each sample, of ClearMap detection was compared against manual counts of c-Fos-immunoreactive (c-Fos-ir) cells in Fiji (Schindelin et al., 2012), yielding 92% precision, 66% recall, and an F1 score of 77% across the cohort (**Figure 2D**).

**Figure 2.**
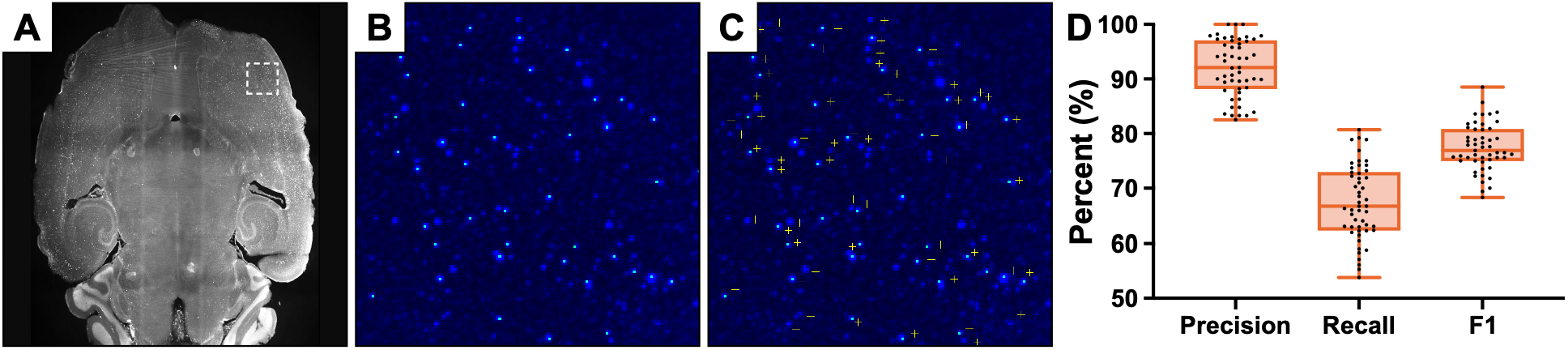
Validation of automated ClearMap detection of c-Fos-immunoreactive cells. Light-sheet fluorescence photomicrographs of whole brain tissue cleared by iDISCO+ processing and labeled with c-Fos-immunoreactive (-ir) signals (**A**) were counted by automated ClearMap (Renier et al., 2016) detection (**B**). Overlap of c-Fos-ir cells detected by manual and automated counting (**C**) determined the precision, which is the percent of cells detected by ClearMap and confirmed manually; recall, which is the percent of cells counted manually and also detected by ClearMap; and overall performance (F1), a percentage measure of cell detection that avoids false positives (**D**).

Registration and object assignment were quality-controlled by overlaying atlas boundaries and detected objects on the autofluorescence and c-Fos volumes and inspecting major anatomical landmarks, including the ventricular boundaries, corpus callosum, external capsule, and major commissures. Detections assigned to fibre tracts or the ventricular system were retained in the exported dataset but were excluded from regional biological interpretation because these assignments were not independently validated. Ventricular assignments may reflect registration or segmentation errors or signal near anatomical boundaries, whereas fibre-tract assignments may also include c-Fos expression in astrocytes, oligodendrocytes, or microglia (Cruz-Mendoza et al., 2022). Regional c-Fos-ir results were reported as both absolute counts and percentages of the total whole-brain count. Absolute counts describe the magnitude of detected c-Fos expression, whereas percentages describe the relative contribution of each region to the brain-wide distribution and minimize sensitivity to between-sample differences in total signal magnitude. Because percentage data are compositional and can obscure global changes, they were interpreted alongside absolute counts.

#### 2.7.2 Plane of section analysis and Nissl-based parcellations

Brightfield images were exported as stitched .tiff files and imported into Affinity Designer 2 (Serif Ltd, Nottingham, UK). Regional boundaries of amygdala sub-nuclei were drawn in a separate layer using the Vector Brush tool in Affinity Designer 2, as previously described (Bono et al., 2022; Negishi et al., 2020). Regional boundaries were identified using cytoarchitectural characteristics, including cell morphology, cell directionality, and Nissl stain intensity, with reference to the *Allen Reference Atlas* (ARA; Dong, 2008). An appropriate ARA level (L) was assigned to each section based on whole-brain white and grey matter patterns.

#### 2.7.3 Manual c-Fos quantification and atlas mapping

In a separate layer, we used the Vector Brush tool in Affinity Designer 2 to place a filled circle on c-Fos-ir cells, then determined the number of circle objects placed within each amygdalar sub-nucleus, as defined by Nissl-based parcellations, to ascertain c-Fos-ir cell counts. All available data points were included to compare the distribution of c-Fos-ir cells across ARA levels. The number of ARA levels contributing to the c-Fos count was held constant across all sleep conditions and sex for each major amygdala division and subnucleus but may vary between divisions or subnuclei. Researchers were blinded to conditions during quantification.

The number of c-Fos-ir cells was corrected for oversampling, as previously described (Bono et al., 2022; Chee et al., 2013; Negishi et al., 2020), using the Abercrombie formula (Abercrombie, 1946): 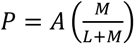 where P is the corrected cell count; A is the original cell count; M is the mean tissue thickness (in μm); and L is the mean cell diameter (in μm). Mean tissue thickness and cell diameter were determined using a ×20 objective *via* fluorescence and brightfield microscopy, respectively. The mean tissue thickness (19.25 μm) was determined from four brain slices. As cell size and morphology may differ between major amygdalar subdivisions, separate correction factors were applaied for each division. Within the cortical subplate, the mean cell diameter (9.97 μm) was determined over 40 cells samples from the basolateral and basomedial amygdala. Within the striatum-like amygdalar nuclei, the mean cell diameter (7.93 μm) was determined over 55 cells samples from the central, medial, and intercalated amygdala.

The relative position of filled circles were transferred to ARA templates (Dong, 2008) in Affinity Designer 2 to produce standardized maps showing representative c-Fos expression across amygdalar levels.

### 2.8 Statistics

#### 2.8.1 iDISCO+

All analyses were performed in Python (version 3.12.9; Python Software Foundation, Wilmington, DE) using the *statsmodels* library for negative binomial regression and custom scripts for data subsetting and multiple comparison correction.

Counts of c-Fos-ir cells were analyzed separately for each brain region using generalized linear models appropriate for overdispersed count data. Specifically, negative binomial regression models with a log link function were fitted to compare c-Fos-ir cell counts between experimental conditions and/or sexes.

For each region, comparisons were restricted to biologically meaningful subsets of the data. Three types of comparisons were performed: (i) sex comparisons within the same sleep condition, (ii) sleep condition comparisons within a single sex, and (iii) sleep condition comparisons pooling both sexes. Subsetting was performed independently for each region to ensure that all models were fit using region specific data only.

Model specification depended on the comparison type. For sex comparisons within a sleep condition, c-Fos-ir cell counts were modeled as a function of sex (value ∼ sex). For sleep condition comparisons within a single sex, counts were modeled as a function of sleep condition (value ∼ sleep condition). For sleep condition comparisons pooling sexes, both group and sex were included as fixed effects (value ∼ sleep condition + sex) to control for sex as a covariate. In all cases, categorical predictors were explicitly ordered so that the first condition served as the reference level. Model coefficients therefore represent log rate ratios comparing the contrasted condition to the reference.

Models were fitted using maximum likelihood estimation. For each comparison and region, the estimated regression coefficient, corresponding rate ratio (exponentiated coefficient), Wald test *p* value, and 95% confidence interval for the rate ratio were extracted. Model fit statistics, including log likelihood and Akaike information criterion (AIC), were also recorded. If a model failed to converge or could not be estimated, the comparison was retained with missing statistics and the fitting error logged.

To control for multiple testing across brain regions, *p* values were adjusted using the Benjamini– Hochberg false discovery rate (FDR) procedure. FDR correction was applied separately for each distinct comparison (e.g., treatment A *vs*. treatment B within males), across all regions with valid *p* values. Adjusted *p* values are reported alongside unadjusted values, and statistical significance was assessed using the FDR corrected results. Plots of fold change per brain region were generated in GraphPad Prism 9.3.1 to visualize region-wise effect sizes against statistical significance. The x-axis represents the estimated log rate ratio and the y-axis −log_10_(FDR). Significance thresholds were highlighted.

#### 2.8.2 Manual c-Fos quantification

In GraphPad Prism 9.3.1, repeated measures three-way ANOVA was used to compare differences in cell counts at major amygdala subdivisions in response to sleep condition and sex. Repeated measures two-way analysis of variance (ANOVA) was used to compare differences in total cell counts from the amygdala between sleep conditions across ARA levels, and to compare differences in total cell counts between sex across ARA levels. Significant main effects and/or interactions proceeded with Bonferroni post-test for pairwise comparisons between sleep conditions or sex.

In SPSS v29 (IBM, Armonk, NY), three-way ANOVA was used to determine main effects of and/or interactions between sex, sleep conditions, and 17 amygdalar subnuclei. Significant main effects and/or interactions were followed-up on with pairwise comparisons of estimated marginal means, maintaining an *α*-level of.05 *via* a Bonferroni correction.

All frequency distribution histograms, bar plots, and lines plots were generated with GraphPad Prism. All data are expressed as the mean ± standard error of the mean (SEM), unless otherwise indicated. Comparisons resulting in *p* < .05 were considered statistically significant.

## 3 Results

### 3.1 Unbiased detection of increased c-Fos expression in the cortical subplate after sleep deprivation

We conducted an unbiased screen for SD-responsive mouse brain structures using tissue clearing and light-sheet microscopy to visualize the immediate early gene c-Fos. c-Fos immunoreactivity was registered to the reference atlas generated by Perens and colleagues (2021; **Figure 1**), and ClearMap (Renier et al., 2016; **Figure 2**) was used to obtain atlas-assigned regional c-Fos-immunoreactive (-ir) cell counts. In line with published whole-brain mapping approaches, absolute regional counts were retained as the measure of activation magnitude. As a complementary analysis of brain-wide distribution, each regional count was also expressed as a percentage of the total c-Fos-ir cells detected in that brain. This within-brain normalization indicates the relative contribution of each structure to the overall c-Fos pattern and reduces sensitivity to global between-sample differences in detection magnitude.

However, because the percentages are compositional and may obscure global changes, they were interpreted alongside absolute counts. Analyses included 13 brain structures: cerebellar cortex, cerebellar nuclei, cortical subplate, hippocampal formation, hypothalamus, isocortex, medulla, midbrain, olfactory areas, pallidum, pons, striatum, thalamus, as well as fiber tracts and ventricular systems. Relative c-Fos measures at rest, after SD, and after RS are reported for male (**Table 3**) and female mice (**Table 4**). As the assignments within fiber tracts and ventricular systems were not validated separately and could not be attributed unequivocally to neuronal activity, they were not interpreted biologically or considered evidence of sleep-deprivation-induced activation within these compartments

**Table 3.** Relative c-Fos expression after sleep deprivation and recovery in male mice.

| Brain structure | Rested<sup>a</sup> (ZT4) |  |  | Sleep deprivation |  |  | Rested<sup>b</sup> (ZT6) |  |  | Recovery sleep |  |  |
|---|---|---|---|---|---|---|---|---|---|---|---|---|
|  | *N* | Mean<sup>c</sup> | SEM | *N* | Mean<sup>c</sup> | SEM | *N* | Mean<sup>c</sup> | SEM | *N* | Mean<sup>c</sup> | SEM |
| CBN | 4 | 0.1 | 0.0 | 4 | 0.1 | 0.1 | 4 | 0.1 | 0.0 | 4 | 0.0 | 0.0 |
| CBX | 4 | 4.4 | 1.1 | 4 | 1.9 | 0.5 | 4 | 2.7 | 0.5 | 4 | 2.0 | 0.3 |
| CTXsp | 4 | 5.1 | 1.3 | 4 | 8.5 | 0.6 | 4 | 6.6 | 1.1 | 4 | 9.1 | 1.3 |
| HPF | 4 | 10.8 | 2.7 | 4 | 12.9 | 1.9 | 4 | 13.4 | 1.9 | 4 | 12.3 | 2.0 |
| HY | 4 | 2.2 | 0.2 | 4 | 3.7 | 0.7 | 4 | 4.6 | 1.0 | 4 | 2.6 | 0.7 |
| ISO | 4 | 31.0 | 5.0 | 4 | 36.4 | 3.5 | 4 | 22.9 | 2.3 | 4 | 35.2 | 5.5 |
| MB | 4 | 7.4 | 0.7 | 4 | 5.3 | 1.3 | 4 | 8.2 | 1.0 | 4 | 6.0 | 1.0 |
| MY | 4 | 2.3 | 0.6 | 4 | 0.8 | 0.1 | 4 | 1.5 | 0.1 | 4 | 1.0 | 0.1 |
| OLF | 4 | 6.6 | 0.6 | 4 | 12.4 | 2.1 | 4 | 5.2 | 0.2 | 4 | 9.8 | 1.4 |
| P | 4 | 2.5 | 0.6 | 4 | 1.3 | 0.3 | 4 | 2.0 | 0.4 | 4 | 1.1 | 0.3 |
| PAL | 4 | 1.0 | 0.2 | 4 | 0.9 | 0.2 | 4 | 1.0 | 0.1 | 4 | 1.2 | 0.4 |
| STR | 4 | 10.6 | 0.8 | 4 | 9.8 | 1.4 | 4 | 15.1 | 2.2 | 4 | 10.9 | 2.6 |
| TH | 4 | 5.2 | 2.5 | 4 | 1.0 | 0.4 | 4 | 4.1 | 1.1 | 4 | 0.9 | 0.2 |
| VS | 4 | 5.0 | 1.0 | 4 | 1.8 | 0.3 | 4 | 6.7 | 1.5 | 4 | 4.2 | 1.1 |
| FT | 4 | 5.8 | 0.8 | 4 | 3.0 | 0.5 | 4 | 5.9 | 0.6 | 4 | 3.7 | 0.2 |

<sup>a</sup> Resting control compared to c-Fos expression following sleep deprivation at Zeitgeber time (ZT) 4
<sup>b</sup> Resting control compared to c-Fos expression following sleep deprivation and recovery sleep at ZT6
<sup>c</sup> Percentage of total c-Fos-immunoreactive cells detected in the whole brain
CBN, cerebellar nuclei; CBX, cerebellar cortex; CTXsp, cortical subplate; FT, fiber tracts; HPF, hippocampal formation; HY; hypothalamus; ISO, isocortex; MB, midbrain; MY, medulla; OLF, olfactory areas; P, pons; PAL, pallidum; SEM, standard error of the mean; STR, striatum; TH, thalamus; VS, ventricular systems

**Table 4.** Relative c-Fos expression after sleep deprivation and recovery in female mice.

| **Brain structure** | **Rested<sup>a</sup> (ZT4)** |  |  | **Sleep deprivation** |  |  | **Rested<sup>b</sup> (ZT6)** |  |  | **Recovery sleep** |  |  |
|---|---|---|---|---|---|---|---|---|---|---|---|---|
|  | ***N*** | **Mean<sup>c</sup>** | **SEM** | ***N*** | **Mean<sup>c</sup>** | **SEM** | ***N*** | **Mean<sup>c</sup>** | **SEM** | ***N*** | **Mean<sup>c</sup>** | **SEM** |
| CBN | 4 | 0.2 | 0.1 | 4 | 0.1 | 0.0 | 4 | 0.0 | 0.0 | 4 | 0.1 | 0.0 |
| CBX | 4 | 4.9 | 1.4 | 4 | 3.7 | 0.8 | 4 | 1.4 | 0.3 | 4 | 2.6 | 0.6 |
| CTXsp | 4 | 4.7 | 0.8 | 4 | 6.3 | 0.5 | 4 | 8.6 | 0.5 | 4 | 8.1 | 0.3 |
| HPF | 4 | 13.6 | 2.1 | 4 | 13.0 | 2.3 | 4 | 11.2 | 1.2 | 4 | 15.9 | 1.9 |
| HY | 4 | 1.8 | 0.5 | 4 | 2.7 | 0.3 | 4 | 3.0 | 0.5 | 4 | 3.0 | 0.8 |
| ISO | 4 | 23.0 | 2.6 | 4 | 26.8 | 5.5 | 4 | 30.5 | 1.1 | 4 | 21.4 | 1.9 |
| MB | 4 | 7.1 | 1.6 | 4 | 7.4 | 0.8 | 4 | 6.7 | 0.7 | 4 | 6.1 | 1.2 |
| MY | 4 | 2.7 | 0.7 | 4 | 1.5 | 0.3 | 4 | 0.8 | 0.1 | 4 | 1.5 | 0.2 |
| OLF | 4 | 4.6 | 1.0 | 4 | 4.6 | 0.5 | 4 | 11.9 | 1.6 | 4 | 7.8 | 1.2 |
| P | 4 | 2.7 | 1.0 | 4 | 1.6 | 0.1 | 4 | 1.1 | 0.4 | 4 | 1.6 | 0.6 |
| PAL | 4 | 0.8 | 0.2 | 4 | 1.0 | 0.2 | 4 | 1.1 | 0.3 | 4 | 1.1 | 0.2 |
| STR | 4 | 16.2 | 3.7 | 4 | 15.9 | 3.7 | 4 | 15.6 | 1.5 | 4 | 18.9 | 3.0 |
| TH | 4 | 3.6 | 2.3 | 4 | 4.3 | 2.0 | 4 | 2.2 | 0.3 | 4 | 3.1 | 1.4 |
| VS | 4 | 7.0 | 0.9 | 4 | 5.5 | 1.2 | 4 | 2.6 | 0.4 | 4 | 4.0 | 0.8 |
| FT | 4 | 7.0 | 1.2 | 4 | 5.7 | 0.3 | 4 | 3.2 | 0.2 | 4 | 4.7 | 0.5 |

<sup>a</sup> Resting control compared to c-Fos expression following sleep deprivation at Zeitgeber time (ZT) 4
<sup>b</sup> Resting control compared to c-Fos expression following sleep deprivation and recovery sleep at ZT6
<sup>c</sup> Percentage of total c-Fos-immunoreactive cells detected in the whole brain
CBN, cerebellar nuclei; CBX, cerebellar cortex; CTXsp, cortical subplate; FT, fiber tracts; HPF, hippocampal formation; HY; hypothalamus; ISO, isocortex; MB, midbrain; MY, medulla; OLF, olfactory areas; P, pons; PAL, pallidum; SEM, standard error of the mean; STR, striatum; TH, thalamus; VS, ventricular systems

Baseline c-Fos expression at ZT4 varied across structures (*F*(14, 180) = 114.7, *p* < .0001). The isocortex, hippocampal formation, and striatum together comprised half of all c-Fos-ir cells detected, while other structures each contributed less than 7% of all c-Fos-ir cells. This baseline distribution of c-Fos-ir cells was comparable between male (**Figure 3A**) and female (**Figure 3B**) mice (*F*(14, 90) = 1.5, *p* = .147) and did not differ between ZT4 and ZT6 in either males (*F*(1, 90) = 5.4e^−13^, *p* > .999) or females (*F*(1, 90) = 6.6e^−19^, *p* > .999).

**Figure 3.**
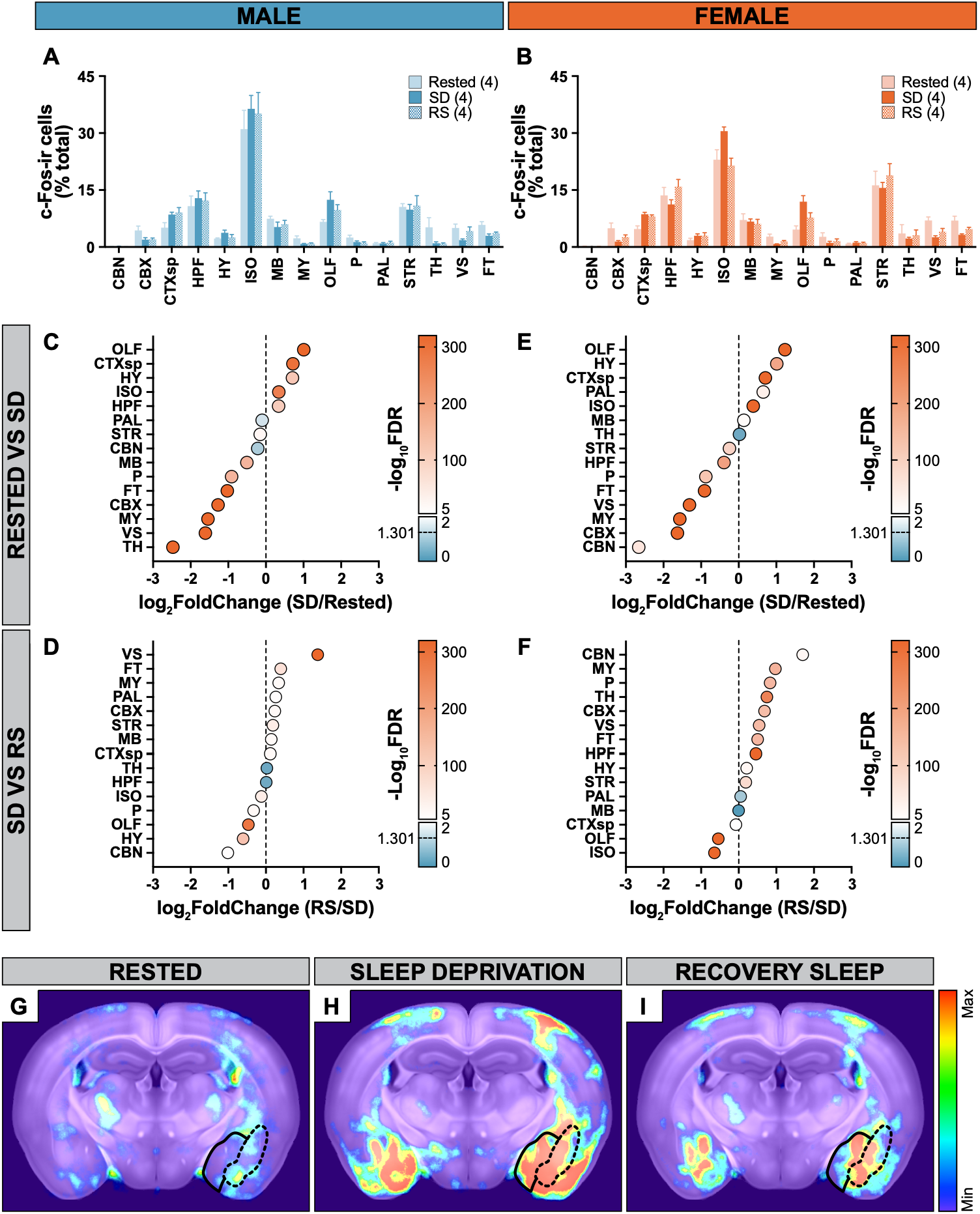
Brain-wide evaluation of c-Fos expression following sleep deprivation. Percentage of c-Fos-immunoreactive (-ir) cells at each high-level structure as a function of total c-Fos-ir cells quantified throughout the whole brain of male (**A**) and female mice (**B**). Data are presented as mean ± SEM. Percent c-Fos expression in each brain structure following sleep deprivation (SD) for 4 h compared to rested male (**C**) or female mice (**E**) or following 2 h of recovery sleep (RS) compared to SD in male (**D**) and female mice (**F**). Log2(fold-change) in percent of c-Fos-ir cells was plotted against −log10(*p*-value) corrected for false discovery rate (FDR). Horizontal dashed line represents *p* = .05. Representative coronal map of (Perens et al., 2021) heat signatures depicting c-Fos-ir cells at rest (**G**), after 4-h SD (**H**), and after SD and 2-h RS (**I**) with solid and dashed black lines defining the striatum-like amygdalar (sAMY) and cortical subplate (CTXsp) division of the amygdala, respectively. CBN, cerebellar nuclei; CBX, cerebellar cortex; CTXsp, cortical subplate; FT, fiber tracts; HPF, hippocampal formation; HY; hypothalamus; ISO, isocortex; MB, midbrain; MY, medulla; OLF, olfactory areas; P, pons; PAL, pallidum; STR, striatum; TH, thalamus; VS, ventricular systems.

Importantly, SD differentially impacted c-Fos expression at each brain structure (*F*(14, 180) = 5.5, *p* = .003), but the structure-specific effect of SD did not differ between sex (*F*(14, 180) = 0.4, *p* < .999). To determine the SD-mediated change in c-Fos expression, we evaluated the fold-change in c-Fos expression and significance of this fold-change at each structure relative to rest (**Table 5**). In males, SD increased c-Fos expression in the olfactory areas, cortical subplate, hypothalamus, isocortex, and hippocampal formation (**Figure 3C**). RS differentially impacted these SD-activated structures, as c-Fos expression reversed in the olfactory areas, hypothalamus, and isocortex; remained elevated in the hippocampal formation; and slightly increased in the cortical subplate (**Figure 3D**). In females, SD increased c-Fos expression in the olfactory areas, hypothalamus, cortical subplate, pallidum, isocortex, and midbrain (**Figure 3E**). Similarly, RS differentially impacted c-Fos expression in these SD-activated structures, as c-Fos expression reversed in the isocortex, olfactory areas, and cortical subplate; remained elevated in the midbrain and pallidum; and further increased in the hypothalamus (**Figure 3F**).

**Table 5.**
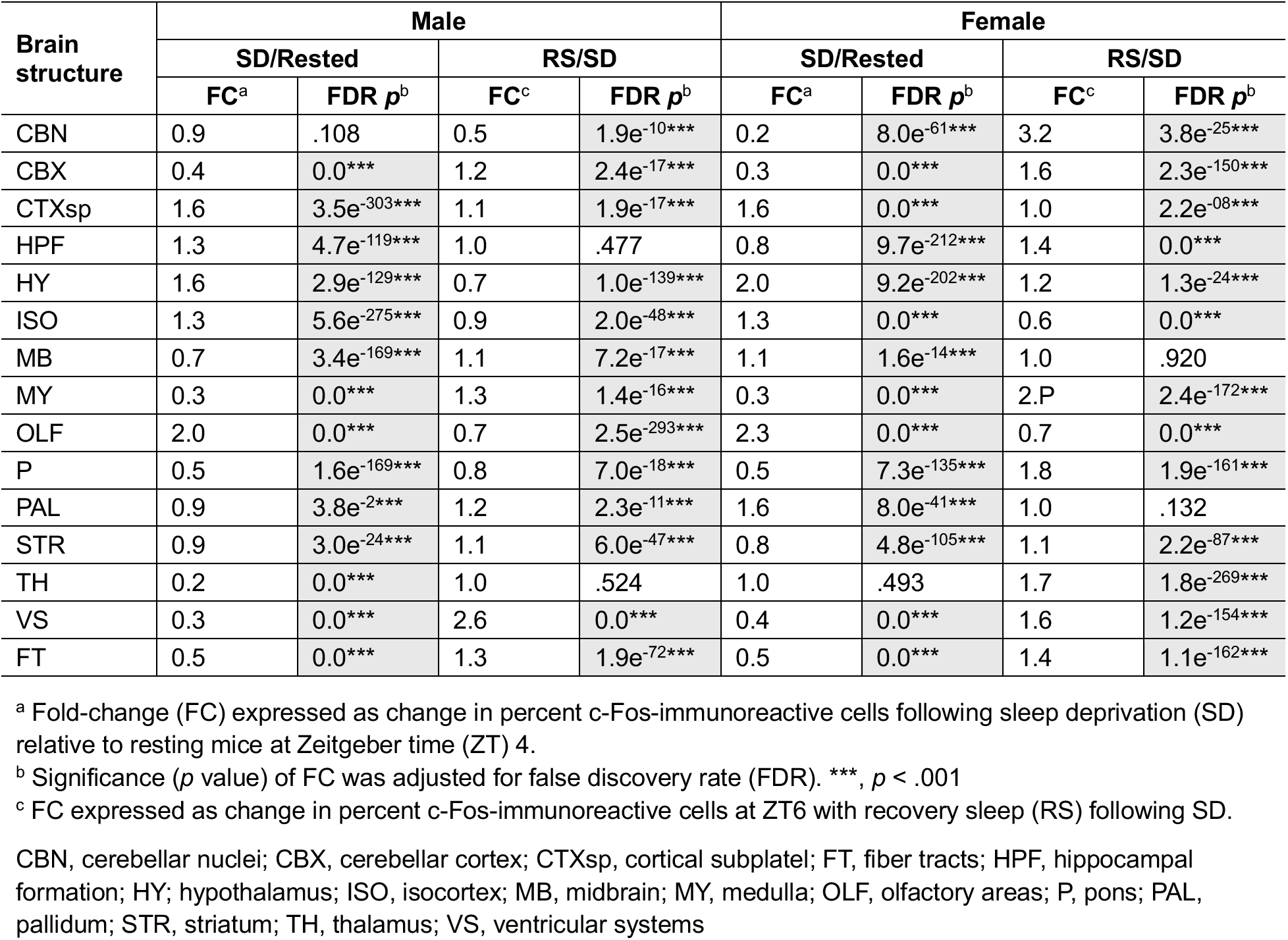
Change in c-Fos expression per brain structure following sleep deprivation and recovery sleep.

| Brain structure | Male |  |  |  | Female |  |  |  |
|---|---|---|---|---|---|---|---|---|
|  | SD/Rested |  | RS/SD |  | SD/Rested |  | RS/SD |  |
|  | FC<sup>a</sup> | FDR ***p***<sup>b</sup> | FC<sup>c</sup> | FDR ***p***<sup>b</sup> | FC<sup>a</sup> | FDR ***p***<sup>b</sup> | FC<sup>c</sup> | FDR ***p***<sup>b</sup> |
| CBN | 0.9 | .108 | 0.5 | 1.9e<sup>-10</sup>*** | 0.2 | 8.0e<sup>-61</sup>*** | 3.2 | 3.8e<sup>-25</sup>*** |
| CBX | 0.4 | 0.0*** | 1.2 | 2.4e<sup>-17</sup>*** | 0.3 | 0.0*** | 1.6 | 2.3e<sup>-150</sup>*** |
| CTXsp | 1.6 | 3.5e<sup>-303</sup>*** | 1.1 | 1.9e<sup>-17</sup>*** | 1.6 | 0.0*** | 1.0 | 2.2e<sup>-08</sup>*** |
| HPF | 1.3 | 4.7e<sup>-119</sup>*** | 1.0 | .477 | 0.8 | 9.7e<sup>-212</sup>*** | 1.4 | 0.0*** |
| HY | 1.6 | 2.9e<sup>-129</sup>*** | 0.7 | 1.0e<sup>-139</sup>*** | 2.0 | 9.2e<sup>-202</sup>*** | 1.2 | 1.3e<sup>-24</sup>*** |
| ISO | 1.3 | 5.6e<sup>-275</sup>*** | 0.9 | 2.0e<sup>-48</sup>*** | 1.3 | 0.0*** | 0.6 | 0.0*** |
| MB | 0.7 | 3.4e<sup>-169</sup>*** | 1.1 | 7.2e<sup>-17</sup>*** | 1.1 | 1.6e<sup>-14</sup>*** | 1.0 | .920 |
| MY | 0.3 | 0.0*** | 1.3 | 1.4e<sup>-16</sup>*** | 0.3 | 0.0*** | 2.P | 2.4e<sup>-172</sup>*** |
| OLF | 2.0 | 0.0*** | 0.7 | 2.5e<sup>-293</sup>*** | 2.3 | 0.0*** | 0.7 | 0.0*** |
| P | 0.5 | 1.6e<sup>-169</sup>*** | 0.8 | 7.0e<sup>-18</sup>*** | 0.5 | 7.3e<sup>-135</sup>*** | 1.8 | 1.9e<sup>-161</sup>*** |
| PAL | 0.9 | 3.8e<sup>-2</sup>*** | 1.2 | 2.3e<sup>-11</sup>*** | 1.6 | 8.0e<sup>-41</sup>*** | 1.0 | .132 |
| STR | 0.9 | 3.0e<sup>-24</sup>*** | 1.1 | 6.0e<sup>-47</sup>*** | 0.8 | 4.8e<sup>-105</sup>*** | 1.1 | 2.2e<sup>-87</sup>*** |
| TH | 0.2 | 0.0*** | 1.0 | .524 | 1.0 | .493 | 1.7 | 1.8e<sup>-269</sup>*** |
| VS | 0.3 | 0.0*** | 2.6 | 0.0*** | 0.4 | 0.0*** | 1.6 | 1.2e<sup>-154</sup>*** |
| FT | 0.5 | 0.0*** | 1.3 | 1.9e<sup>-72</sup>*** | 0.5 | 0.0*** | 1.4 | 1.1e<sup>-162</sup>*** |

<sup>a</sup> Fold-change (FC) expressed as change in percent c-Fos-immunoreactive cells following sleep deprivation (SD) relative to resting mice at Zeitgeber time (ZT) 4.
<sup>b</sup> Significance (*p* value) of FC was adjusted for false discovery rate (FDR). \*\*\*, *p* < .001
<sup>c</sup> FC expressed as change in percent c-Fos-immunoreactive cells at ZT6 with recovery sleep (RS) following SD.
CBN, cerebellar nuclei; CBX, cerebellar cortex; CTXsp, cortical subplatel; FT, fiber tracts; HPF, hippocampal formation; HY; hypothalamus; ISO, isocortex; MB, midbrain; MY, medulla; OLF, olfactory areas; P, pons; PAL, pallidum; STR, striatum; TH, thalamus; VS, ventricular systems

Together, these analyses show that the largest SD-induced increases in c-Fos expression occurred in the cortical subplate and olfactory areas of both male and female mice. Since activity in the olfactory areas may reflect experimenter interaction during SD, we focused on c-Fos differences within the cortical subplate at rest (**Figure 3G**), after SD (**Figure 3H**), and after RS (**Figure 3I**). The cortical subplate includes the claustrum, endopiriform nuclei, and some amygdalar regions, but qualitative assessment of heat signatures reflecting c-Fos-ir cells within the cortical subplate after SD was localized to areas within the amygdala rather than the claustrum or endopiriform nuclei (**Figure 3H**). Therefore, we focussed on resolving c-Fos expression patterns within the amygdala complex.

### 3.2 SD increased c-Fos expression throughout the male but not female amygdala

To determine whether SD uniformly or differentially activated each amygdalar subnucleus, we performed plane-of-section analyses to parcellate these subnuclei, then generated high resolution spatial maps to compare the distribution of c-Fos-ir cells across experimental groups (**Figure 4**). To increase spatial resolution, we first quantified c-Fos-ir cells at ∼150 μm intervals rostrocaudally within the amygdala of wildtype male and female mice at rest, SD, and RS. The mouse amygdala comprises two major divisions: the medially located striatum-like amygdalar nuclei (sAMY) and the laterally positioned cortical subplate (CTXsp; **Figure 5A**). It begins at ARA level L55 (−0.08 mm from Bregma (β)) and ends at L86 (β −3.28 mm), with the largest designated area towards the middle (L65–L75; **Figure 5A2, 5A3**).

**Figure 4.**
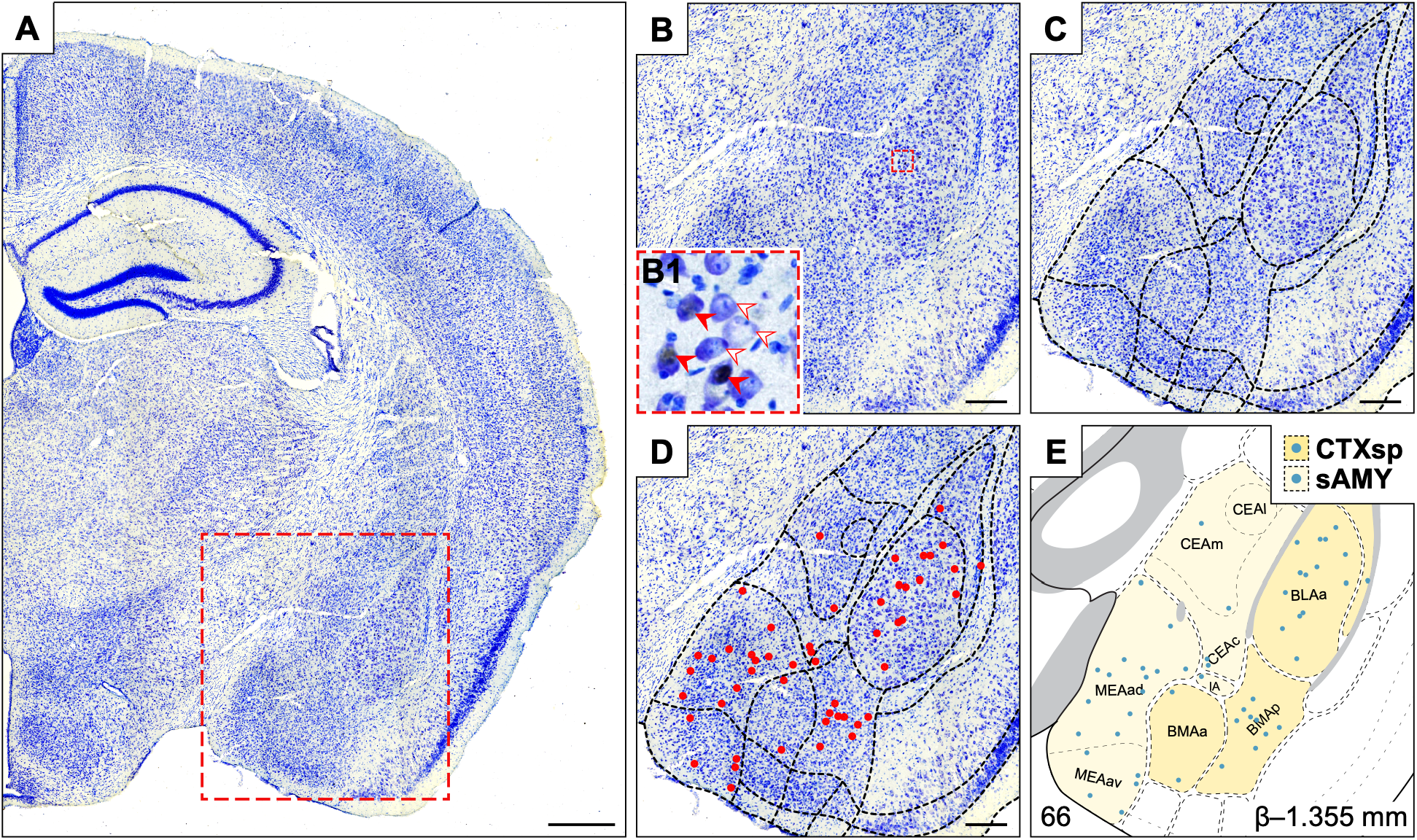
Schematic of c-Fos quantification and spatial mapping in the mouse amygdala. Brightfield photomicrograph of Nissl-stained coronal brain section co-labelled for c-Fos immunoreactivity (**A**) with red dashed box indicating the region of interest shown in **B**. High magnification photomicrograph (**B1**) from the dashed region outlined in **B** showing Nissl-stained blue cell body (open arrowhead) expressing c-Fos immunoreactivity with black nucleus (filled arrowhead). Amygdalar subnuclei were parcellated (dashed black lines) according to the *Allen Reference Atlas* (ARA; Dong, 2008; **C**) and c-Fos-immunoreactive cells were marked with a red circle (**D**). The red circles were then mapped to the corresponding ARA (Dong, 2008) template (**E**). Scale bar: 500 μm (**A**); 200 μm (**B**–**D**).

**Figure 5.**
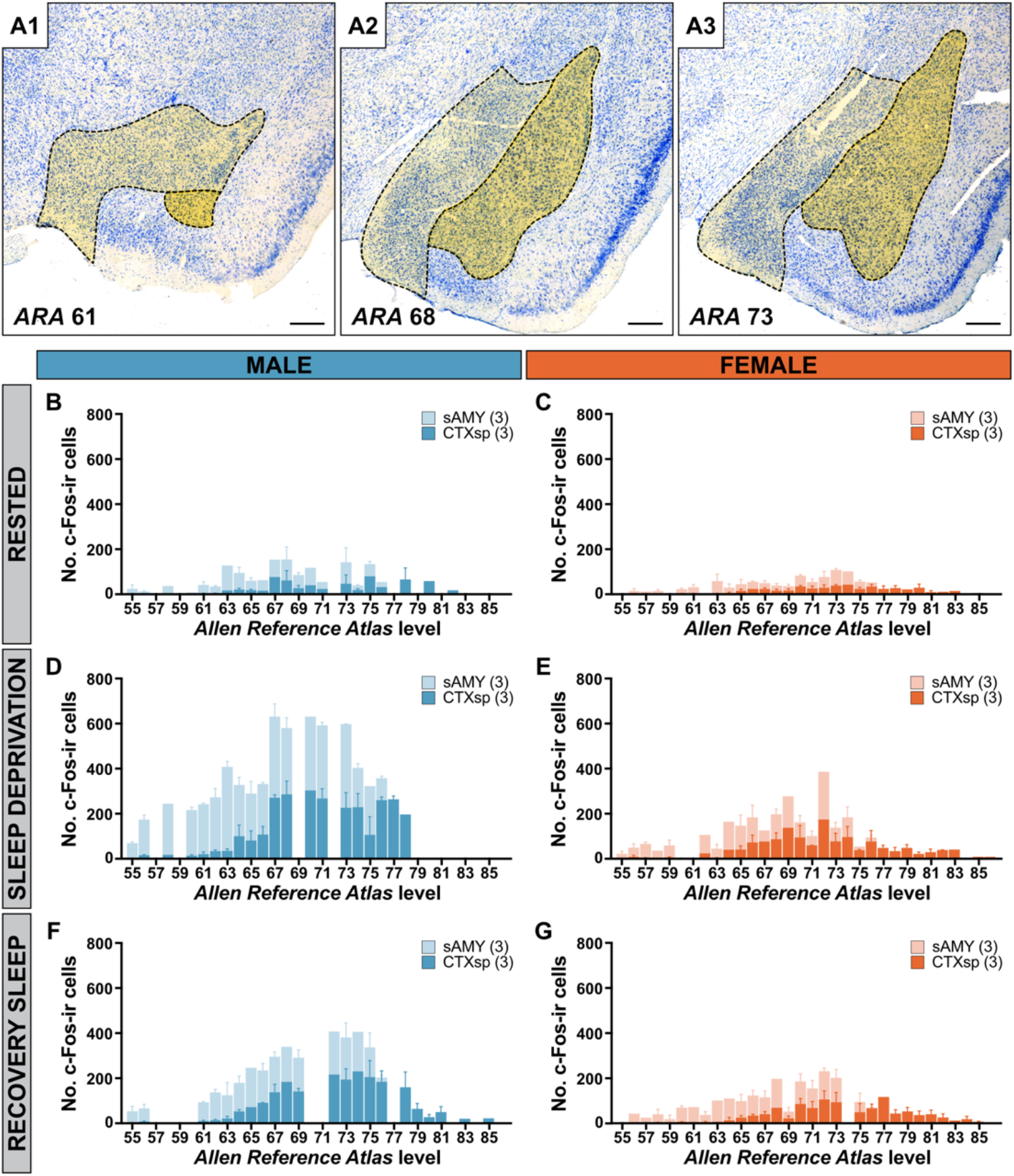
Sleep deprivation increased c-Fos expression rostrocaudally in male but not female amygdala. Brightfield photomicrographs of c-Fos-immunoreactive (-ir) and Nissl-stained cells within the anterior (**1**), middle (**2**), and posterior (**3**) amygdala overlayed with a light and dark yellow shade to define the striatum-like amygdalar (sAMY) and cortical subplate (CTXsp) division of the amygdala, respectively (**A**). Distribution of c-Fos-ir cells within the sAMY (light shading) and CTXsp (dark shading) at each *Allen Reference Atlas* (ARA; Dong, 2008) level of the amygdala in male (blue) and female (orange) mice at rest (**B**, **C**), after 4-h sleep deprivation (SD; **D**, **E**), and SD with 2-h recovery sleep (RS; **F**, **G**). Missing bars indicate that tissue sections were not captured at that ARA and were unavailable for analysis. Data are presented as mean ± SEM where data are available from at least two brains; absence of error bars indicate cell counts from one brain. Scale bar: 250 μm (**A1**–**3**)

Baseline c-Fos expression counts did not vary across ARA levels (*F*(20, 80) = 1.6, *p* = .083) in the male (**Figure 5B**) or female amygdala (**Figure 5C**), and c-Fos distribution across ARA levels also did not differ between sex (*F*(20, 80) = 0.8, *p* = .762).

Following SD, c-Fos expression increased across ARA levels and reached statistical significance in male (*F*(18, 72) = 2.0, *p* = .023; **Figure 5D**) but not female mice (*F*(27, 108) = 1.0, *p* = .444; **Figure 5E**). Indeed, c-Fos expression following SD differed between sex (*F*(1, 4) = 20.3, *p* = .011). In male mice, sleep opportunity after SD tended to lower c-Fos expression (*F*(1, 4) = 7.5, *p* = .0.052; **Figure 5F**) compared to SD and resembled resting levels (*F*(1, 4) = 4.8, *p* = .094). c-Fos expression after RS in male mice were comparable to that in the female amygdala, as there were no differences in c-Fos expression between sexes after RS (*F*(1, 4) = 3.1, *p* = .151; **Figure 5G**).

The sAMY and CTXsp amygdalar divisions have varying prominence across ARA levels and may overlap at some ARA levels (**Figure 6A–C**). We found that despite these anatomical aspects, c-Fos expression did not differ by division (*F*(1, 24) = 1.4, *p* = .243). Additionally, baseline c-Fos expression was comparable between the sAMY and CTXsp of male (*p* > .999) and of female (*p* > .999) mice (**Figure 6A, 6D**).

**Figure 6.**
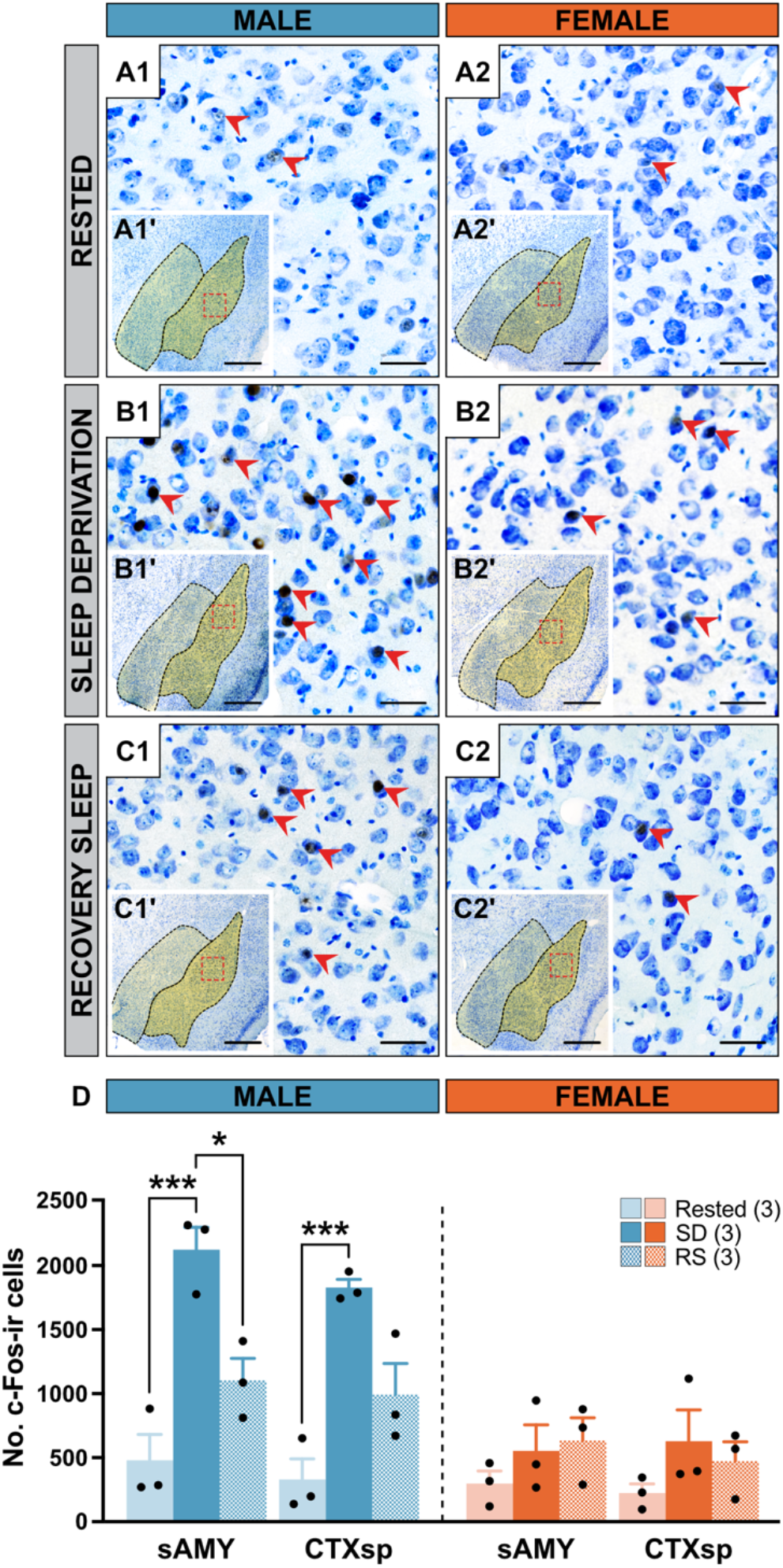
Sleep deprivation increased c-Fos expression at both amygdala divisions of male but not female mice. High magnification brightfield photomicrographs of c-Fos-immunoreactive (-ir) at Nissl-stained cells (red arrowheads) within the amygdala of male (**1**) and female (**2**) mice at rest (**A**), after 4-h sleep deprivation (SD; **B**), and SD with 2-h recovery sleep (RS; **C**). Red dashed box in **A**–**C1’** and **A**–**C2’** indicates region shown at high magnification in **A**–**C1** and **A**–**C2**, respectively. Overlayed yellow shading in **A**–**C1’** and **A**–**C2’** define the striatum-like amygdalar (sAMY, light shade) and cortical subplate (CTXsp, dark shade) division of the amygdala. Number of c-Fos-ir cells within the sAMY and CTXsp divisions of the male (blue) and female (orange) amygdala at rest (light shade), after SD (dark shade), and RS (checkered; **D**). Data are presented as mean ± SEM. Three-way repeated measures ANOVA with Bonferroni’s post-hoc test, where: **, *p* < .01; ***, *p* < .001. Scale bar: 500 μm (**A**–**C1, A**–**C2**); 100 μm (**A**–**C1’, A**–**C2’**).

Importantly, c-Fos expression differed by sleep condition in a sex-dependent manner (*F*(2, 24) = 13.6, *p* < .001), but sleep condition similarly affected the sAMY and CTXsp (*F*(2, 24) = .01, *p* = .993). Specifically, SD increased the number of c-Fos-ir cells in the sAMY (*p* < .001) and CTXsp (*p* < .001) of male mice, but both divisions did not reach statistically significant increases in female mice (sAMY: *p* > .999; CTXsp: *p* > .999; **Figure 6B**, **6D**). RS differentially impacted the sAMY and CTXsp (**Figure 6C**). c-Fos expression within the sAMY of male mice decreased from SD (*p* = .023) to baseline levels (*p* > .999), while that in the CTXsp trended towards baseline levels (*p* = .824) but was not significantly reduced from SD expression (*p* = .151). In females, c-Fos expression in the sAMY and CTXsp following RS remained similar to baseline (sAMY: *p* > .999; CTXsp: *p* > .999) and SD (sAMY: *p* > .999; CTXsp: *p* > .999; **Figure 6D**).

Overall, these analyses across ARA levels and amygdalar divisions indicated broad SD-induced c-Fos expression within the male but not female amygdala.

### 3.3 Specificity in SD-induced c-Fos by amygdalar subnuclei

Since different subnuclei within the sAMY and CTXsp may respond uniquely to sleep loss, we next assessed c-Fos expression across experimental groups within each subnuclei (**Figure 7**). Additionally, we also considered whether SD spatially defined c-Fos distribution within each subnucleus of the male (**Figure 8**) and female amygdala (**Figure 9**). We first quantified the number of c-Fos-ir cells at each subnucleus and then mapped the spatial distribution of these cells onto ARA templates to generate standardized neuroanatomical maps (Dong, 2008).

**Figure 7.**
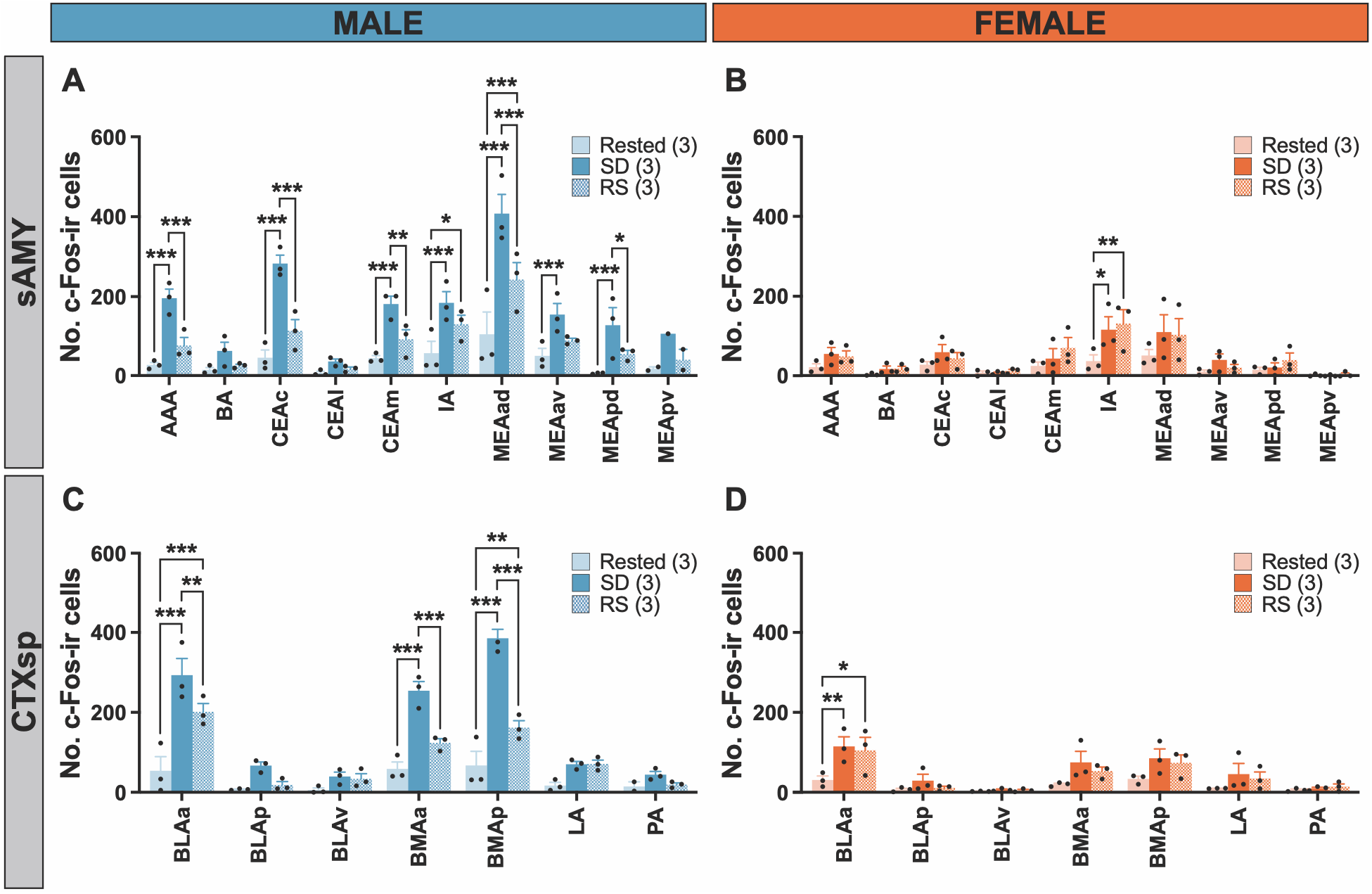
Sleep deprivation increased c-Fos expression at select amygdalar subnuclei. Number of c-Fos-immunoreactive (-ir) cells quantified at individual amygdalar subnuclei within the striatum-like amygdala (sAMY; **A**, **B**) and cortical subplate (CTXsp; **C**, **D**) of male (blue) and female (orange) mice at rest (light shade), after 4-h sleep deprivation (SD; dark shade), and SD with 2-h recovery sleep (RS; checkered). Number of *Allen Reference Atlas* levels included for each amygdala subnucleus was standardized. Data are presented as mean ± SEM. Three-way repeated measures ANOVA with Bonferroni’s post-hoc test, where: *, *p* < .05; **, *p* < .01; ***, *p* < 0.001. AAA, anterior amygdalar area; BA, bed nucleus of the accessory olfactory tract; BLAa, anterior part, BLAp, posterior part, BLAv, ventral part of basolateral amygdalar nucleus; BMAa, anterior part, BMAp, posterior part of basomedial amygdalar nucleus; CEAc, capsular part, CEAl, lateral part, CEAm, medial part of central amygdalar nucleus; IA, intercalated amygdalar nucleus; LA, lateral amygdalar nucleus; MEAad, anterodorsal part, MEAav, anteroventral part, MEApd, posterodorsal part, MEApv, posteroventral part of medial amygdalar nucleus; PA, posterior amygdalar nucleus.

**Figure 8.**
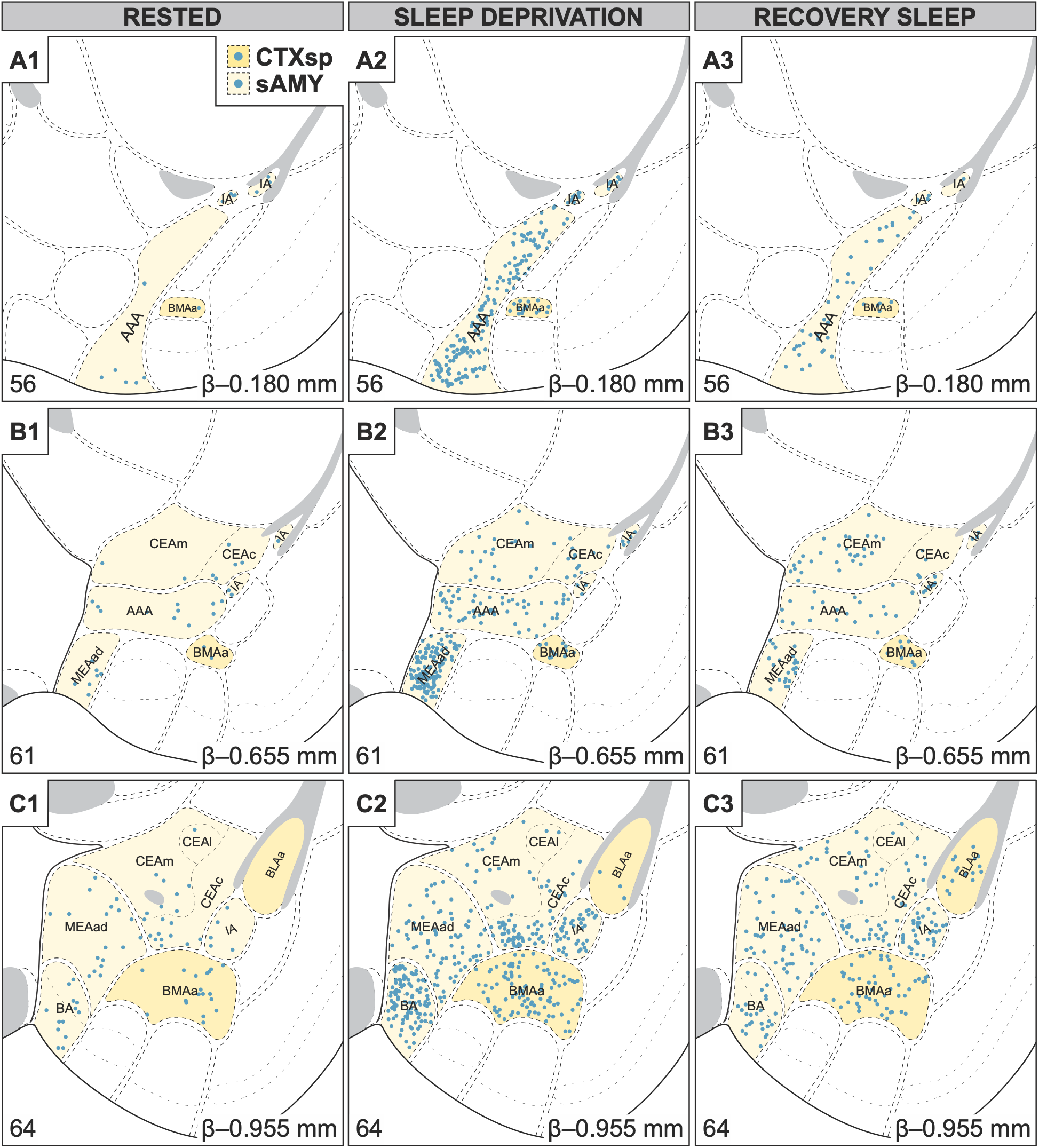
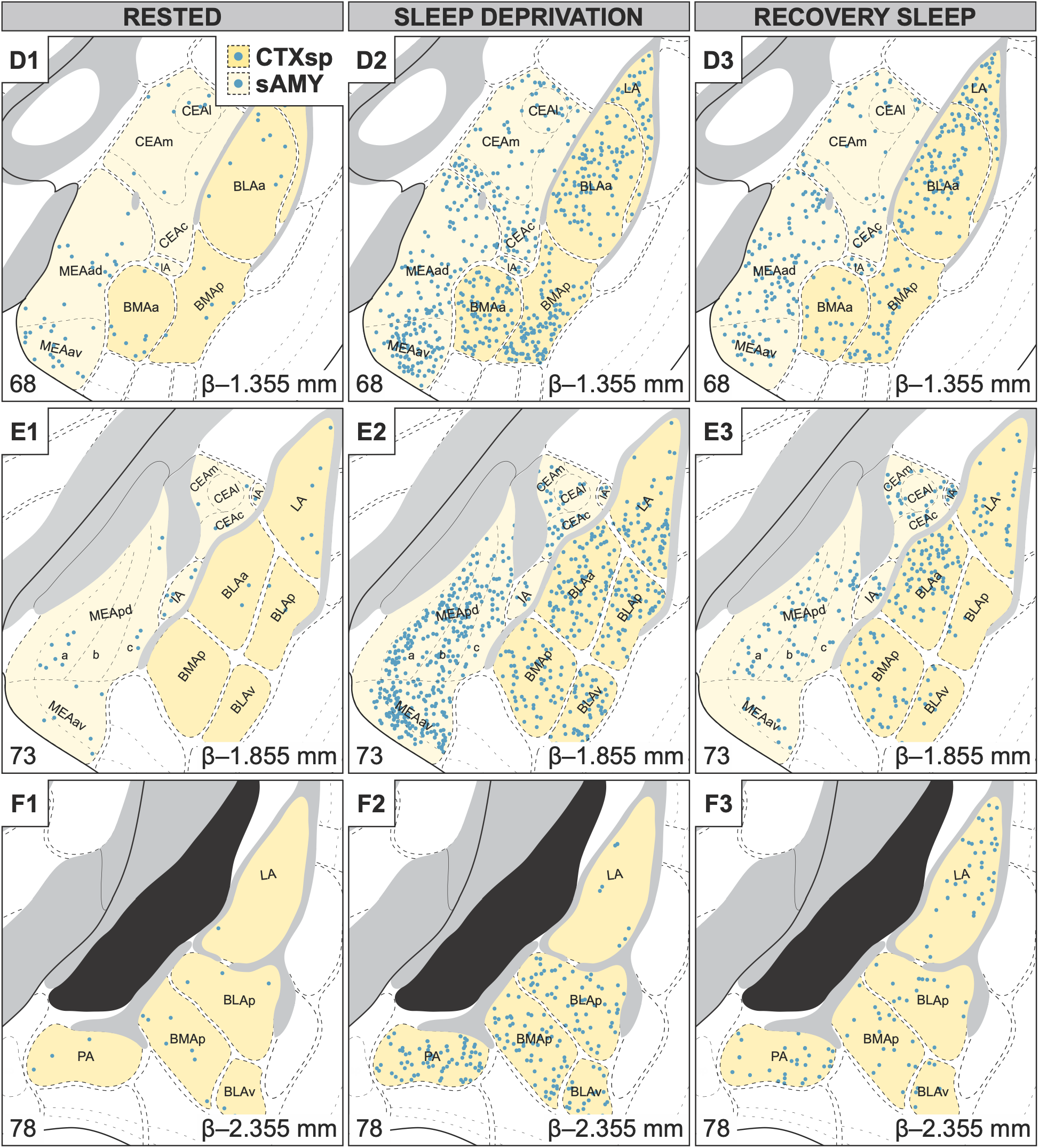
Differential c-Fos distribution in the amygdala of male mice with sleep loss. Representative, unilateral coronal atlas maps from the *Allen Reference Atlas* (ARA; Dong, 2008) arranged rostrocaudally (**A**–**F**) showing the relative distribution of c-Fos-immunoreactive cells in the male mouse amygdala at rest (**1**), after 4-h sleep deprivation (**2**), and after 2-h recovery sleep (**3**). Subnuclei comprising the striatum-like amygdala (sAMY) and cortical subplate (CTXsp) divisions are shaded in light yellow and dark yellow, respectively. Each panel includes the corresponding ARA level (bottom right), Bregma level (β; top right), and subnuclei labels using formal ARA nomenclature.

**Figure 9.**
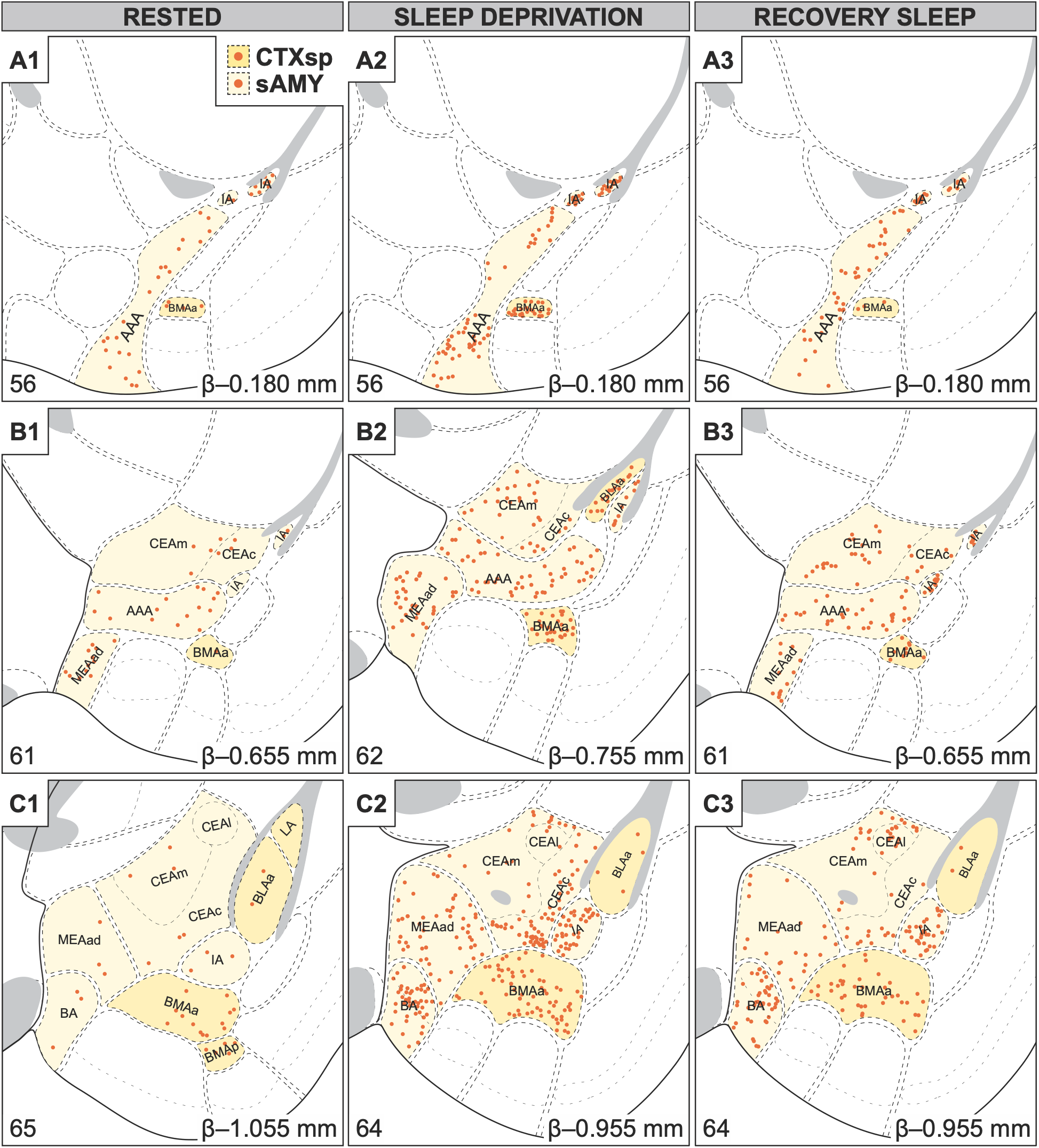
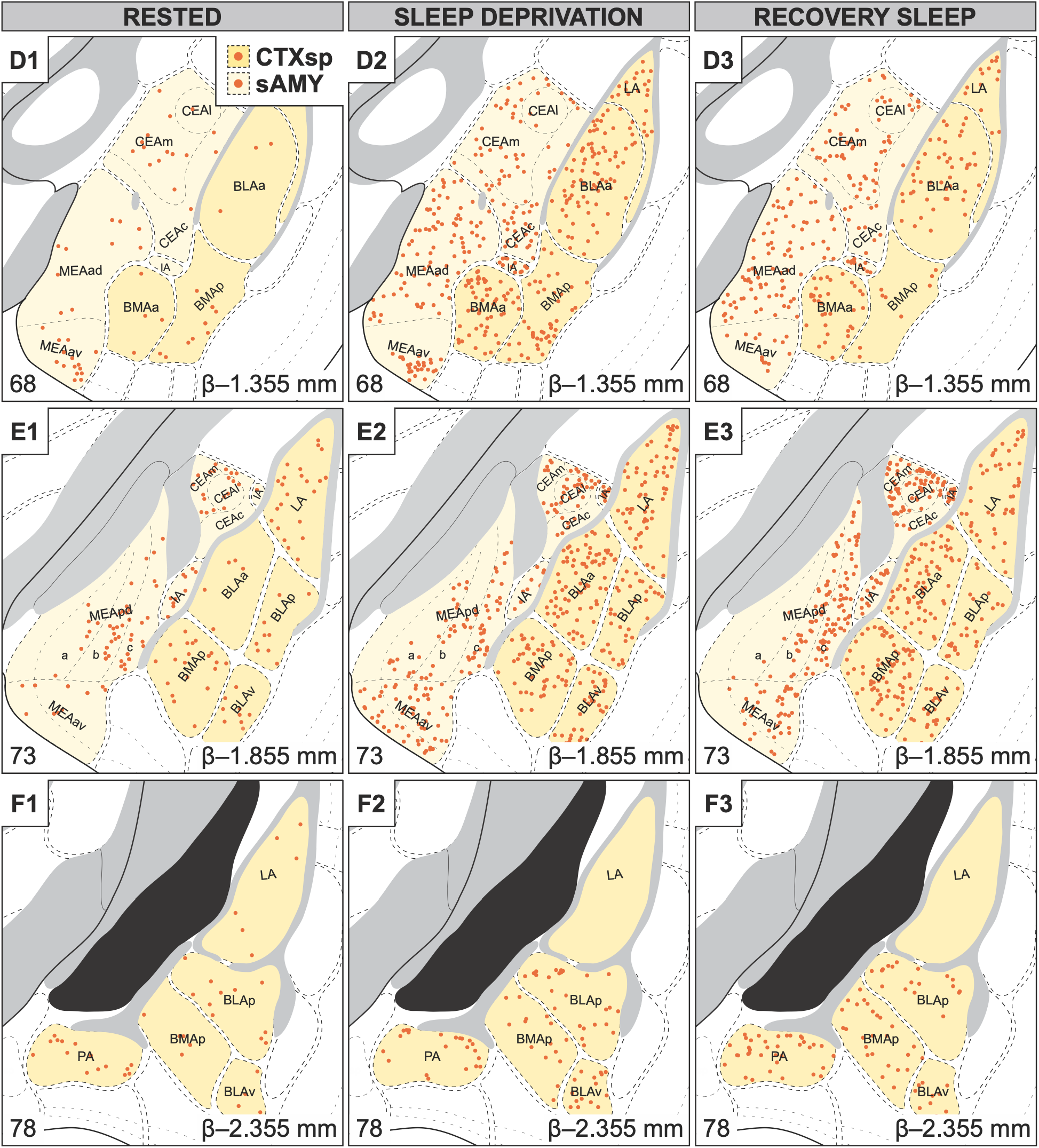
Differential c-Fos distribution in the amygdala of female mice with sleep loss. Representative, unilateral coronal atlas maps from the *Allen Reference Atlas* (ARA; Dong, 2008) arranged rostrocaudally (**A**–**F**) showing the relative distribution of c-Fos-immunoreactive cells in the female mouse amygdala at rest (**1**), after 4-h sleep deprivation (**2**), and after 2-h recovery sleep (**3**). Subnuclei comprising the striatum-like amygdala (sAMY) and cortical subplate (CTXsp) divisions are shaded in light yellow and dark yellow, respectively. Each panel includes the corresponding ARA level (bottom right), Bregma level (β; top right), and subnuclei labels using formal ARA nomenclature.

Overall, c-Fos expression varied significantly between sex (*F*(1, 199) = 2291.1, *p* < .001), among subnuclei (*F*(16, 199) = 34.8, *p* < .001), and in response to sleep condition (*F*(2, 199) = 141.2, *p* < .001). Importantly, c-Fos expression differed by sleep condition in a sex-(*F*(2, 199) = 60.1, *p* < .001) and subnucleus-dependent manner (*F*(32, 199) = 4.6, *p* < .001). We thus assessed differences in c-Fos expression due to sleep condition with pairwise comparisons at individual subnuclei within the CTXsp and sAMY. The results of these comparisons are listed for males in **Table 6** and for females in **Table 7**.

**Table 6.** Pairwise comparisons at individual amygdalar subnuclei for each sleep condition in male mice.

| **Amygdalar subnucleus** | **Rested** |  |  | **Sleep deprivation** |  |  | **Recovery sleep** |  |  | ***p*** |  |  |
|---|---|---|---|---|---|---|---|---|---|---|---|---|
|  | ***N*** | **Mean** | **SEM** | ***N*** | **Mean** | **SEM** | ***N*** | **Mean** | **SEM** | **Rested *vs.* SD** | **SD *vs.* RS** | **Rested *vs.* RS** |
| **Striatum-like amygdalar nuclei** |  |  |  |  |  |  |  |  |  |  |  |  |
| AAA | 3 | 29.0 | 6.1 | 3 | 196.0 | 22.9 | 3 | 77.7 | 20.2 | < .001*** | < .001*** | .252 |
| BA | 3 | 16.7 | 5.8 | 3 | 64.0 | 22.0 | 3 | 27.3 | 3.5 | .278 | .577 | 1.000 |
| CEAc | 3 | 47.0 | 19.4 | 3 | 282.7 | 21.1 | 3 | 114.3 | 28.0 | < .001*** | < .001*** | .052 |
| CEAl | 3 | 9.0 | 4.0 | 3 | 37.7 | 6.2 | 3 | 19.3 | 4.4 | .923 | 1.000 | 1.000 |
| CEAm | 3 | 45.7 | 6.7 | 3 | 181.3 | 19.7 | 3 | 93.3 | 23.9 | < .001*** | .006** | .272 |
| IA | 3 | 58.3 | 29.8 | 3 | 184.3 | 28.3 | 3 | 131.0 | 22.3 | < .001*** | .176 | .031* |
| MEAad | 3 | 105.7 | 55.7 | 3 | 407.7 | 48.3 | 3 | 242.7 | 42.6 | < .001*** | < .001*** | < .001*** |
| MEAav | 3 | 51.7 | 18.8 | 3 | 155.0 | 28.0 | 3 | 90.3 | 5.5 | < .001*** | .066 | .508 |
| MEApd | 3 | 8.0 | 1.0 | 3 | 128.3 | 44.1 | 3 | 56.7 | 8.9 | < .001*** | .034* | .252 |
| MEApv | 2 | 19.5 | 4.5 | 1 | 107.0 | N/A | 2 | 42.0 | 26.0 | .116 | .371 | 1.000 |
| **Cortical subplate** |  |  |  |  |  |  |  |  |  |  |  |  |
| BLAa | 3 | 53.7 | 35.6 | 3 | 293.0 | 41.7 | 3 | 201.3 | 20.7 | < .001*** | .004** | < .001*** |
| BLAp | 3 | 7.0 | 1.2 | 3 | 66.7 | 9.1 | 3 | 18.3 | 8.4 | .104 | .259 | >.999 |
| BLAv | 3 | 5.7 | 5.2 | 3 | 39.3 | 11.1 | 3 | 33.7 | 13.0 | .693 | >.999 | .957 |
| BMAa | 3 | 58.3 | 17.3 | 3 | 254.0 | 23.1 | 3 | 123.7 | 10.4 | < .001*** | < .001*** | .062 |
| BMAp | 3 | 67.0 | 35.5 | 3 | 385.3 | 22.4 | 3 | 161.7 | 17.5 | < .001*** | < .001*** | .003** |
| LA | 3 | 16.7 | 8.0 | 3 | 70.0 | 7.0 | 3 | 71.0 | 9.5 | .176 | >.999 | .162 |
| PA | 2 | 14.5 | 11.5 | 3 | 44.0 | 8.3 | 3 | 19.0 | 4.4 | >.999 | >.999 | >.999 |

\*, *p* < .05; \*\*, *p* < .01; \*\*\*, *p* < .001
AAA, anterior amygdalar area; BA, bed nucleus of the accessory olfactory tract; BLAa, anterior part, BLAp, posterior part, BLAv, ventral part of basolateral amygdalar nucleus; BMAa, anterior part, BMAp, posterior part of basomedial amygdalar nucleus; CEAc, capsular part, CEAl, lateral part, CEAm, medial part of central amygdalar nucleus; IA, intercalated amygdalar nucleus; LA, lateral amygdalar nucleus; MEAad, anterodorsal part, MEAav, anteroventral part, MEApd, posterodorsal part, MEApv, posteroventral part of medial amygdalar nucleus;  PA, posterior amygdalar nucleus; RS, recovery sleep; SEM, standard error of the mean; SD, sleep deprivation

**Table 7.**
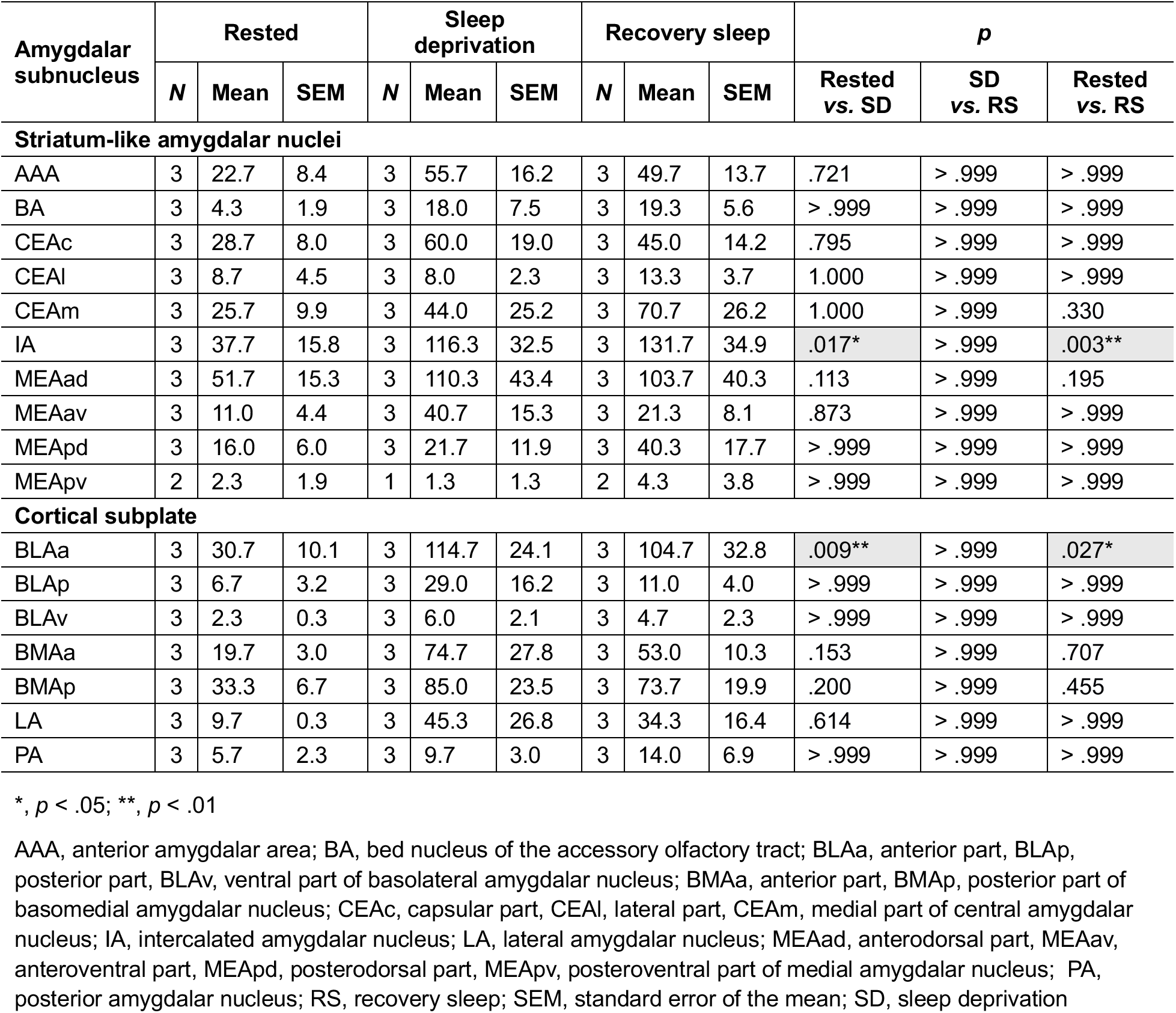
Pairwise comparisons at individual amygdalar subnuclei for each sleep condition in female mice.

| **Amygdalar subnucleus** | **Rested** |  |  | **Sleep deprivation** |  |  | **Recovery sleep** |  |  | ***p*** |  |  |
|---|---|---|---|---|---|---|---|---|---|---|---|---|
|  | ***N*** | **Mean** | **SEM** | ***N*** | **Mean** | **SEM** | ***N*** | **Mean** | **SEM** | **Rested *vs.* SD** | **SD *vs.* RS** | **Rested *vs.* RS** |
| **Striatum-like amygdalar nuclei** |  |  |  |  |  |  |  |  |  |  |  |  |
| AAA | 3 | 22.7 | 8.4 | 3 | 55.7 | 16.2 | 3 | 49.7 | 13.7 | .721 | >.999 | >.999 |
| BA | 3 | 4.3 | 1.9 | 3 | 18.0 | 7.5 | 3 | 19.3 | 5.6 | >.999 | >.999 | >.999 |
| CEAc | 3 | 28.7 | 8.0 | 3 | 60.0 | 19.0 | 3 | 45.0 | 14.2 | .795 | >.999 | >.999 |
| CEAl | 3 | 8.7 | 4.5 | 3 | 8.0 | 2.3 | 3 | 13.3 | 3.7 | 1.000 | >.999 | >.999 |
| CEAm | 3 | 25.7 | 9.9 | 3 | 44.0 | 25.2 | 3 | 70.7 | 26.2 | 1.000 | >.999 | .330 |
| IA | 3 | 37.7 | 15.8 | 3 | 116.3 | 32.5 | 3 | 131.7 | 34.9 | .017* | >.999 | .003** |
| MEAad | 3 | 51.7 | 15.3 | 3 | 110.3 | 43.4 | 3 | 103.7 | 40.3 | .113 | >.999 | .195 |
| MEAav | 3 | 11.0 | 4.4 | 3 | 40.7 | 15.3 | 3 | 21.3 | 8.1 | .873 | >.999 | >.999 |
| MEApd | 3 | 16.0 | 6.0 | 3 | 21.7 | 11.9 | 3 | 40.3 | 17.7 | >.999 | >.999 | >.999 |
| MEApv | 2 | 2.3 | 1.9 | 1 | 1.3 | 1.3 | 2 | 4.3 | 3.8 | >.999 | >.999 | >.999 |
| **Cortical subplate** |  |  |  |  |  |  |  |  |  |  |  |  |
| BLAa | 3 | 30.7 | 10.1 | 3 | 114.7 | 24.1 | 3 | 104.7 | 32.8 | .009** | >.999 | .027* |
| BLAp | 3 | 6.7 | 3.2 | 3 | 29.0 | 16.2 | 3 | 11.0 | 4.0 | >.999 | >.999 | >.999 |
| BLAv | 3 | 2.3 | 0.3 | 3 | 6.0 | 2.1 | 3 | 4.7 | 2.3 | >.999 | >.999 | >.999 |
| BMAa | 3 | 19.7 | 3.0 | 3 | 74.7 | 27.8 | 3 | 53.0 | 10.3 | .153 | >.999 | .707 |
| BMAp | 3 | 33.3 | 6.7 | 3 | 85.0 | 23.5 | 3 | 73.7 | 19.9 | .200 | >.999 | .455 |
| LA | 3 | 9.7 | 0.3 | 3 | 45.3 | 26.8 | 3 | 34.3 | 16.4 | .614 | >.999 | >.999 |
| PA | 3 | 5.7 | 2.3 | 3 | 9.7 | 3.0 | 3 | 14.0 | 6.9 | >.999 | >.999 | >.999 |

\*, *p* < .05; \*\*, *p* < .01
AAA, anterior amygdalar area; BA, bed nucleus of the accessory olfactory tract; BLAa, anterior part, BLAp, posterior part, BLAv, ventral part of basolateral amygdalar nucleus; BMAa, anterior part, BMAp, posterior part of basomedial amygdalar nucleus; CEAc, capsular part, CEAl, lateral part, CEAm, medial part of central amygdalar nucleus; IA, intercalated amygdalar nucleus; LA, lateral amygdalar nucleus; MEAad, anterodorsal part, MEAav, anteroventral part, MEApd, posterodorsal part, MEApv, posteroventral part of medial amygdalar nucleus;  PA, posterior amygdalar nucleus; RS, recovery sleep; SEM, standard error of the mean; SD, sleep deprivation

#### 3.3.1 sAMY: SD activated the intercalated amygdalar nucleus in both sexes

In males, SD significantly increased c-Fos expression in six subnuclei: anterior amygdalar area (AAA); capsular (CEAc) and medial (CEAm) parts of the central amygdalar nucleus; intercalated amygdalar nucleus (IA); and anterodorsal (MEAad), anteroventral (MEAav), and posterodorsal (MEApd) parts of the medial amygdalar nucleus (**Figure 7A**). In females, SD significantly increased c-Fos expression in the IA only (**Figure 7B**).

##### Anterior amygdalar area (AAA)

c-Fos expression in the male AAA significantly increased following SD and returned towards baseline levels with RS (**Figure 7A**). These effects were not observed in females (**Figure 7B**).

The distribution of SD-induced c-Fos-ir cells was relatively even throughout the male AAA and more dense anteriorly (**Figure 8A**) than posteriorly (**Figure 8B**). In the female AAA, c-Fos-ir cells after SD avoided the center anteriorly (**Figure 9A**) but was evenly distributed posteriorly (**Figure 9B**).

##### Bed nucleus of the accessory olfactory tract (BA)

SD did not significantly increase c-Fos expression in the BA of either male (**Figure 7A**) or female mice (**Figure 7B**). However, c-Fos-ir cells present after SD appeared uniformly distributed in males (**Figure 8C**) yet clustered dorsally in females (**Figure 9C**). c-Fos expression remained unchanged following RS relative to baseline.

##### Central amygdalar nucleus (CEA)

In males, c-Fos expression after SD significantly increased within the medial (CEAm) and capsular (CEAc) part of the CEA and returned towards baseline with RS (**Figure 7A**). Interestingly, this sensitivity to SD did not affect the lateral part of the CEA (CEAI). In females, SD did not alter c-Fos expression in any part of the CEA (**Figure 7B**). c-Fos-ir cells following SD were most prevalent ventrally in the CEA, especially in the male (**Figure 8B**–**D**) and female CEAc (**Figure 9B**–**D**).

##### Intercalated amygdalar nucleus (IA)

In both male (**Figure 7A**) and female mice (**Figure 7B**), SD increased c-Fos expression in the IA, and this SD-induced c-Fos expression remained elevated even after RS. “Intercalated,” by definition, refers to being inserted between layers, thus the IA is an amorphous structure. In effect, no clear patterns emerged in the distribution of c-Fos-ir cells in the IA (**Figure 8A**–**E**, **Figure 9A**–**E**).

##### Medial amygdalar nucleus (MEA)

**In male mice,** SD increased c-Fos expression in the anterodorsal (MEAad), anteroventral (MEAav), and posterodorsal parts of the MEA (MEApd; **Figure 7A**). While c-Fos expression also appeared elevated in the posteroventral MEA (MEApv), this did not reach statistical significance (**Figure 7A**). Each MEA subnucleus responded differently to RS, as c-Fos expression decreased but not to baseline levels in the

MEAad, remained elevated in the MEAav, and decreased to baseline levels in the MEApd (**Figure 7A**).

With respect to distribution in males, SD-induced c-Fos-ir cells in the MEAad were densely and uniformly distributed in anterior sections then became sparser and favoured the ventral or dorsal borders posteriorly (**Figure 8B**–**D**). Within the MEAav, c-Fos-ir cells were densely clustered dorsally throughout (**Figure 8C**–**E**). c-Fos cells in the MEApd were evenly distributed in sublayer-a, -b, and -c though there were fewer cells in sublayer-c (**Figure 8E**). c-Fos cells in the MEApv were more densely clustered in the ventral aspect of the structure (data not shown).

In females, none of the MEA subnuclei showed significant changes after SD or RS (**Figure 7B**). Nevertheless, we did observe patterns in c-Fos-ir cell distribution after SD. Cells in the MEAad were uniformly dispersed (**Figure 9B**–**D**). Within the MEAav, SD-induced c-Fos-ir cells clustered ventrolaterally in the anterior (**Figure 9D**) but were more uniformly distributed posteriorly (**Figure 9E**). In contrast to males, most cells in the MEApd appeared evenly distributed in sublayer-c (**Figure 9E**). Only five c-Fos cells were detected in the female MEApv and so no pattern could be discerned (data not shown).

#### 3.3.2 CTXsp: SD activated the anterior basolateral amygdala in both sexes

In males, SD increased c-Fos expression in three subnuclei: anterior part of the basolateral amygdalar nucleus (BLAa), and anterior (BMAa) and posterior (BMAp) parts of the basomedial amygdalar nucleus **(Figure 7C)**. In females, only the BLAa showed significantly increased c-Fos expression after SD (**Figure 7D**).

##### Basolateral amygdala (BLA)

SD increased c-Fos expression in the anterior BLA (BLAa) in male (**Figure 7C**) and female mice (**Figure 7D**) but that in the posterior (BLAp) or ventral BLA (BLAv) were unaffected. After RS, c-Fos expression partially reduced in males (**Figure 7C**), but persisted in the female BLAa (**Figure 7D**).

The BLAa is the longest structure but coincides with the BLAp and BLAv posteriorly, and the distribution of c-Fos-ir cells in the BLAa was similar in male (**Figure 8C**–**E**) and female mice (**Figure 9C**–**E**). The rostral BLAa structure was sparsely labeled with c-Fos-ir cells (**Figure 8C**; **Figure 9C**), but as the BLAa expands (e.g., towards L68), c-Fos-ir cells featured towards the medial aspect of the BLAa (**Figure 8D**; **Figure 9D**) and then were uniformly distributed in caudal sections (**Figure 8E**; **Figure 9E**).

The distribution of c-Fos-ir cells were consistent between the male and female BLAp and BLAv. c-Fos-ir cells were uniformly distributed within these structures in the male (**Figure 8E**, **8F**) and female (**Figure 9E**, **9F**), but we did not detect a significant increase in c-Fos expression in these structures (**Figure 7C**, **7D**).

##### Basomedial amygdalar nucleus (BMA)

In male mice, c-Fos expression increased in both the anterior BMA (BMAa) and posterior BMA (BMAp) after SD (**Figure 7C**). After RS, c-Fos expression fully returned to resting levels in the BMAa but only partially reversed in the BMAp (**Figure 7C**). Neither part of the BMA significantly responded to SD in female mice (**Figure 7D**).

In males, c-Fos-ir cells were distributed uniformly across the rostracaudal extent of the BMAa under all conditions (**Figure 8A**–**D**). In the BMAp, c-Fos-ir cells clustered ventrally in anterior sections (**Figure 8D**) but were evenly distributed posteriorly (**Figure 8E**, **8F**). Similar distribution patterns were observed in the BMAa (**Figure 9C**, **9D**) and BMAp (**Figure 9D**–**F**) of female mice.

##### Lateral amygdala (LA)

We observed very few c-Fos cells in the resting LA of male or female mice, and c-Fos expression in the LA did not increase following SD or RS of either male (**Figure 7C**) or female mice (**Figure 7D**).

Following SD, LA c-Fos-ir cells appeared anteriorly in male (**Figure 8D**, **8E**) and female mice (**Figure 9D**, **9E**) and were devoid of cells posteriorly (**Figure 8F**, **9F**). In the anterior LA, c-Fos cells appeared dorsally in both sexes (**Figure 8D**, **9D**) but as the LA progressed posteriorly, c-Fos cells appeared ventrally in the males (**Figure 8E**) and dorsally in the females (**Figure 9F**).

##### Posterior amygdala (PA)

c-Fos expression did not increase in SD or RS in the PA of male (**Figure 7C**) or female mice (**Figure 7D**). c-Fos cells that appeared following SD tended to be ventrally distributed in males (**Figure 8F**) and those appearing in females favored along the dorsal PA border (**Figure 9F**).

Taken together, the male amygdala showed broad c-Fos recruitment following SD, and this expression can be seen at several amygdalar subnuclei. In the female amygdala, c-Fos expression was highly selective and appeared at the BLAa and IA only. Interestingly, while SD-induced c-Fos expression typically reversed with RS in male mice, any increased c-Fos expression seen in female mice persisted even with RS.

## 4 Discussion

In addition to established roles in emotion and mood regulation, the amygdala also has emergent roles in sleep regulation. Therefore, it is a promising candidate region linking mood disruptions following sleep loss. Nevertheless, the specific amygdalar subnuclei activated following SD were previously unresolved. We screened the intact mouse brain for SD-activated structures using c-Fos expression as an activity marker and identified prominent increases in c-Fos in the cortical subplate after SD. As the cortical subplate largely comprises amygdalar subnuclei, we resolved SD-induced c-Fos expression at each amygdalar subnucleus rostrocaudally. Interestingly, SD broadly increased c-Fos expression in the entire male amygdala but restricted c-Fos increase to only the BLA and IA of the female amygdala.

### Unique activation of cortical subplate following sleep deprivation

Relative to the whole brain, SD produced the greatest and most significant fold-change increase in c-Fos expression in the olfactory areas, cortical subplate, isocortex, and hypothalamus of both sexes. This is consistent with previous studies showing regionally heterogeneous responses to SD in the mouse forebrain (Thompson et al., 2010). Furthermore, our brain-wide c-Fos patterns are consistent with Fos-dependent activity in TRAP2 reporter mice after six hours of SD (Urushihata et al., 2023). As the hypothalamus is known to be SD-activated (Alam et al., 2014; Basheer et al., 1997; Gong et al., 2004; Vyazovskiy et al., 2009), the isocortex contributes to sleep homeostasis (Gerashchenko et al., 2008; Gong et al., 2004; Krone et al., 2021; Morairty et al., 2013; Sherin et al., 1996), and olfactory activity may relate to the experimental design to initiate and maintain SD (Guthrie et al., 1993; Sorge et al., 2014), we focused on the evoked changes within the cortical subplate.

The cortical sublate (CTXsp) presented as a novel and compelling SD-activated structure from whole brain analyses. The CTXsp comprises the claustrum, endopiriform nucleus, and some amygdalar subnuclei (Dong, 2008), but SD-induced c-Fos in the CTXsp localized to the amygdala only. The amygdala also includes subnuclei in the striatum, but as striatum-like amygdala (sAMY) regions are only a small part of the striatum, any change in the sAMY may have been masked in our top-level screen. We thus parcellated each CTXsp and sAMY amygdalar subnucleus rostrocaudally to confirm SD-induced amygdalar activation. In males, SD increased c-Fos expression in the BLAa and BMA of the CTXsp but nearly all sAMY subnuclei increased c-Fos expression following SD. By contrast, SD-mediated c-Fos expression was restricted in the female amygdala and observed in only one CTXsp (i.e., BLAa) and sAMY subnucleus (i.e., IA).

### Blunted amygdalar response to sleep deprivation in females

SD induced greater increases in amygdalar c-Fos expression in males than females. Consistent with previous studies showing SD-induced c-Fos expression in the amygdala of male mice (Thompson et al., 2010) and rats (Cirelli et al., 1995; Semba et al., 2001), we confirmed that SD increased c-Fos expression in the male amygdala and resolved that this response was specific to subnuclei within both the sAMY and CTXsp. However, SD did not alter c-Fos expression from resting levels in the female amygdala.

Few studies have examined SD-induced brain activation specifically in females. In ovariectomized female rats, a 6-h SD increased c-Fos expression across arousal-promoting, limbic, and neuroendocrine regions (Deurveilher et al., 2008). Other work also indicated that while baseline c-Fos expression in vigilance states may vary between phases of the estrous cycle, the homeostatic response to SD appears unaffected (Schwierin et al., 1998). In our study, SD tended to increase amygdalar c-Fos expression in females, but the effect was smaller than in males and did not reach statistical significance. This attenuated response may reflect lower sensitivity or tighter regulation of activity-dependent signaling during acute SD. Alternatively, it may arise from sex differences in circuit recruitment, sleep pressure, stress responsivity, hormonal state, or c-Fos kinetics.

This preclinical observation is broadly consistent with reported sex differences in objective sleep phenotypes, although direct functional testing is required to determine whether females are more resilient to SD. Despite reporting greater difficulty initiating sleep and poorer perceived sleep quality (Lindberg et al., 1997; Middelkoop et al., 1996), women generally have shorter polysomnographic sleep latency, longer total sleep time, and a greater proportion of slow-wave sleep than men (Bixler et al., 2009; Goel et al., 2005; Redline et al., 2004; Vitiello et al., 2004). Women also show greater low-frequency electroencephalographic (EEG) power during habitual sleep (Dijk et al., 1989). However, slow wave activity primarily indexes sleep intensity and homeostatic sleep pressure and does not, by itself, indicate more efficient recovery. In mice, sex-dependent differences in baseline sleep architecture have also been reported (Paul et al., 2006), whereas homeostatic recovery following a 6-h SD was broadly comparable between males and females (Koehl et al., 2006). Therefore, the attenuated amygdalar response in females may reflect sex-dependent neural recruitment during acute sleep loss but cannot be interpreted as physiological resilience without direct measures of sleep architecture, homeostatic rebound, and functional recovery.

### Amygdalar activation was subnucleus specific

Although SD did not broadly increase amygdalar c-Fos expression in female mice, it selectively increased activation in the BLAa and IA, two subnuclei that were also recruited in males.

Previous studies also indicate that BLA recruitment depends on the duration and context of sleep loss. An acute 6-h SD increased BLA c-Fos expression (Cirelli et al., 1995) but not from a sustained 24-h SD (Montes-Rodríguez et al., 2019; Xu et al., 2026). The BLAa also emerged as a SD-responsive region using targeted recombination in active populations 2 (TRAP2)-labeled cells, although the number of TRAP2-labelled cells in the BLA decreased following a 6-h SD (Urushihata et al., 2023). This difference may reflect the distinct temporal windows captured by the two approaches. Conventional c-Fos immunoreactivity tags proteins 90 min after induction, providing a relatively immediate endpoint measure, whereas TRAP2 requires activation by a tamoxifen analogue to ensure recombination. This means that the Fos-dependent activity could be captured over a broader window that may extend beyond the intended SD period, unintentionally overlapping with sleep onset, which we show can rapidly reverse c-Fos expression. Therefore, the observation that BLA is sensitive to SD remains consistent across studies, while the direction and magnitude of change appears context-dependent.

The functional contribution of SD-induced BLAa activation is yet unclear. One possibility is that the BLA serves a homeostatic sleep function, as different BLA populations can regulate REM (Hasegawa et al., 2022, 2024; Machida et al., 2021) and NREM states (Zhu et al., 1998).

Another possibility pertains to the more canonical functions of the BLA in regulating fear and anxiety, and assigning emotional valence (Janak & Tye, 2015). Notably, human imaging studies report increased global amygdalar reactivity to negative emotional stimuli after SD (Yoo et al., 2007) and of specifically the left BLA during fear acquisition (Feng et al., 2023). Thus, BLAa activation during SD could contribute to heightened emotionality.

Within the sAMY, the IA was similarly, and significantly, activated by SD in both sexes. The IA comprises densely packed inhibitory cells that cluster along the white matter tracts encapsulating the LA and BLA (Millhouse, 1986; Nitecka & Ben-Ari, 1987). Their primary role is to regulate signal transmission within the amygdala, such as by gating the flow of sensory information from the BLA to the CEA (Amano et al., 2010; Paré et al., 2003; Royer et al., 1999). IA cells are important for fear extinction (Likhtik et al., 2008) and social function (Kuerbitz et al., 2018), but little is known about them in the context of sleep beyond maintaining a constant firing rate across sleep-wake states (Collins & Paré, 1999). The increase in c-Fos expression at the IA following SD may be related to concurrent BLA activation. As the BLA can excite IA cells (Royer et al., 1999; Royer & Paré, 2002), and IA cells can inhibit the BLA (Asede et al., 2015; Mańko et al., 2011), coincident activation of the IA and BLA may reflect feedforward inhibition during SD. Alternatively, increased IA activity may gate SD-related BLAa output to other amygdalar structures.

#### Broad amygdalar subnuclei activation in male mice

The male amygdala exhibited broad activation after SD. In addition to the BLAa and IA, SD also increased c-Fos expression in the anterior amygdalar area (AAA) and within subdivisions of the central (CEA), medial (MEA), and basomedial (BMA) amygdala. Existing literature has shown that the CEA (Xu et al., 2026), MEA (Deurveilher et al., 2008), and BMA (Cirelli et al., 1995) are SD-activated. The AAA lacks clear anatomical boundaries (Müller & O’Rahilly, 2006; Swanson, 2018, Supporting Table C), and its function remains poorly understood, with no reported role in sleep to date. However, it does receive direct input from the main olfactory bulb (Hintiryan et al., 2012), thus AAA activation may be related to experimenter interaction during SD.

SD-activated CEA cells may be responding to SD in a homeostatic or stress-related manner. Different neurochemical CEA populations are directly implicated in sleep regulation, including neurotensin cells that promote NREM sleep (Ma et al., 2019) and somatostatin cells that promote arousal in the presence of acute stress (Yao et al., 2025). Activation in the CEA may also be species-dependent, as it was not seen in rat models (Kalinchuk et al., 2010; Mashoodh et al., 2008).

The BMA and MEA are candidate regions through which SD-associated activity could influence affect-related processing. The BMA is a subnucleus that encodes information about aversive contexts to supress fear and anxiety (Adhikari et al., 2015; Rajbhandari et al., 2021). Thus, SD-induced BMA activation may be to contextualize and respond emotionally to sleep loss.

Emerging evidence also indicates that the MEA plays an important role in integrating sleep and anxiety (Li et al., 2024), for example at the posteroventral part of the MEA. However, we did not detect a significant increase in c-Fos expression in this MEA subdivision. Alternatively, as the MEA is well-known for managing innate social behaviours (Prakash et al., 2025) like aggression (Unger et al., 2015), it is possible that MEA activation reflected a defensive reaction to the experimenter during SD. Furthermore, as male rodents exhibit more aggression and territoriality than female counterparts, this may explain why MEA activation appeared in males only.

#### Brain-wide and amygdalar response to recovery sleep

The olfactory areas, isocortex, hypothalamus, and cortical subplate showed distinct patterns of c-Fos expression following the recovery period. The largest decrease occurred in the olfactory areas of both sexes. Although this reversal is consistent with removal of sensory and experimenter-related stimulation present during SD (Urushihata et al., 2023), the present design cannot determine the specific contribution of these factors. RS decreased c-Fos expression in the isocortex, and this decrease was greater and more significant in females. This is consistent with previous reports that wake-associated cortical c-Fos expression declines with RS (Basheer et al., 1997; Vyazovskiy et al., 2009), but specific cortical populations such as cortical cells expressing neuronal nitric oxide synthase (nNOS) are preferentially recruited during RS (Gerashchenko et al., 2008; Morairty et al., 2013). Together, these findings suggest that cortical c-Fos dynamics across SD and RS are not uniform and are cell-type dependent.

In contrast, c-Fos expression in the hypothalamus and cortical subplate remained elevated following RS. However, persistent c-Fos expression alone cannot confirm continued neuronal activity or a direct role in sleep maintenance, particularly because sleep was not physiologically quantified during SD or RS, and sustained c-Fos signal may reflect residual SD-induced c-Fos.

Nevertheless, independent studies have demonstrated recruitment of specific hypothalamic neuronal populations during sleep and following SD (Alam et al., 2014; Gong et al., 2004). The apparent decrease in c-Fos expression observed in the male hypothalamus should also be interpreted cautiously because tissue damage associated with mounting for light-sheet microscopy may compromise c-Fos quantification in this region. In addition, we also detected a decrease in c-Fos expression in the female cortical subplate following RS, but this fold-change was less than 1% of SD levels.

Within the amygdala, c-Fos expression remained elevated in the IA and BLAa following the recovery period, although expression in the male BLAa showed a partial return toward rested levels. This pattern is particularly interesting in relation to Bocchio et al. (2016) who identified a population of nNOS-expressing GABAergic neurons situated along the boundaries of the BLA that showed c-Fos recruitment during undisturbed sleep following SD, but not during SD itself. The population identified here appears spatially and temporally distinct. These cells appeared in the medial aspect of the BLAa and were evident following both SD and RS. These findings therefore support heterogeneity among neuronal populations recruited across SD and subsequent recovery within and around the BLA, although the neurochemical identity of the c-Fos-ir cells identified here remains unknown.

Independent functional evidence further indicates that the BLA can modulate sleep-related network activity. Optogenetic manipulation of BLA glutamatergic neurons during spontaneous REM bidirectionally altered REM-associated hippocampal EEG theta activity without altering overall sleep amount or propensity (Machida et al., 2021). Thus, rather than demonstrating that persistent c-Fos expression in the BLAa reflects a direct role in maintaining RS, the present findings point to the BLAa as a candidate region recruited during SD and the subsequent recovery period, that could participate in sleep-related network processes. Similarly, persistent c-Fos expression in the IA may reflect continued recruitment of circuitry interacting with the BLA, but the present data do not establish the direction or functional significance of this interaction.

#### Theoretical and methodological considerations

As the amygdala contributes greatly to assessing and responding to stressful stimuli, the c-Fos activation may, in part, be due to stress from the SD process or experimenter handling. This could account for the decrease in c-Fos during RS in regions such as the AAA, CEA, and MEA in male mice. Additionally, the CEA and MEA are known to modulate the hypothalamic-pituitary axis (Herman et al., 2020; Smith & Vale, 2006). SD activates these subnuclei, and their reduced activity during RS may support maintaining alertness during a stressful period. In contrast, the sustained activation in the BLAa in RS could be more important for sleep-related outcomes.

Our SD protocol, while commonly used in the literature (e.g., Matsumoto et al., 2020; Rolls et al., 2015), may not have captured SD-sensitive structures equally in both sexes. We monitored the animals for 4 h and employed gentle handling when animals showed signs of sleepiness (e.g., closing eyes, hunched back). This was sufficient to induce c-Fos expression broadly in male mice but only selectively in female mice. Although this more restricted response may reflect reduced sensitivity to sleep loss, it could alternatively indicate slower recruitment of the female amygdala, with longer SD (e.g., 6- or 8-h) required to achieve activation comparable to that observed in males. Thus, the sex difference may reflect either a higher threshold for sleep-loss-induced amygdala activation or distinct temporal dynamics of c-Fos recruitment in females.

Finally, c-Fos expression in some subnuclei remained elevated following recovery sleep, but it is not clear whether this reflected their contribution to sleep maintenance, insufficient recovery sleep, or residual SD-related activity. Furthermore, as the animals were not monitored during RS, and we did not record their sleep-wake architecture, we cannot confirm the amount or extent of recovery sleep each animal received.

## Conclusion

Whole-brain screening identified prominent SD-related increase in c-Fos expression in the amygdalar structures of the cortical subplate. Extending high-resolution neuroanatomical analyses across the entire amygdala revealed sex-dependent recruitment of amygdalar subnuclei following SD. In males, SD induced robust c-Fos expression across multiple subnuclei, whereas in females, significant increases were restricted to the BLAa and IA. These sex differences were also evident during recovery sleep (RS), as SD-induced c-Fos expression readily returned toward baseline in the male amygdala, whereas increases observed in females remained elevated following RS. Together these findings demonstrated that acute sleep loss recruits amygdalar subcircuits in a region-and sex-dependent manner but do not yet establish that the more restricted response in females reflects greater resilience to SD. Future studies integrating physiological sleep measures with cell type-specific neurochemical, electrophysiological, and behavioural analyses will be required to determine their functional significance and potential contributions to sleep-loss-induced deficits in affective and mood regulation.

## Conflicts of interest

The authors have no conflicts of interest to declare.

## Data availability

The data that support the findings of this study are available from the corresponding author upon reasonable request.

## Notes

### Competing Interest Statement

The authors have declared no competing interest.

### Funding

- European Research Council
- Natural Sciences and Engineering Research Council

